# Clamping, bending, and twisting inter-domain motions in the misfold-recognising portion of UDP-glucose:glycoprotein glucosyl-transferase

**DOI:** 10.1101/2019.12.25.888438

**Authors:** Carlos P. Modenutti, Juan I. Blanco Capurro, Roberta Ibba, Snežana Vasiljević, Mario Hensen, Dominic S. Alonzi, Anu V. Chandran, Johan C. Hill, Jonathan Rushton, Abhinav Kumar, Simone Rubichi, Andrea Lia, Gábor Tax, Lucia Marti, Angelo Santino, Marcelo A. Martí, Nicole Zitzmann, Pietro Roversi

**Author notes:** These authors contributed equally to this work (Carlos P. Modenutti, Juan I. Blanco Capurro, Roberta Ibba).

## Abstract

UDP-glucose:glycoprotein glucosyltransferase (UGGT) is the only known glycoprotein folding quality control checkpoint in the eukaryotic glycoprotein secretory pathway. When the enzyme detects a misfolded glycoprotein in the Endoplasmic Reticulum (ER), it dispatches it for ER retention by re-glucosylating it on one of its N-linked glycans. Recent crystal structures of a fungal UGGT have suggested the enzyme is conformationally mobile. Here, a negative stain electron microscopy reconstruction of UGGT in complex with a monoclonal antibody confirms that the misfold-sensing N-terminal portion of UGGT and its C-terminal catalytic domain are tightly associated. Molecular Dynamics (MD) simulations capture UGGT in so far unobserved conformational states, giving new insights into the molecule’s flexibility. Principal component analysis of the MD trajectories affords a description of UGGT’s overall inter-domain motions, highlighting three types of inter-domain movements: bending, twisting and clamping. These inter-domain motions modify the accessible surface area of the enzyme’s central saddle, likely enabling the protein to recognize and re-glucosylate substrates of different sizes and shapes, and/or re-glucosylate N-linked glycans situated at variable distances from the site of misfold. We propose to name “Parodi limit” the maximum distance between a site of misfolding on a UGGT glycoprotein substrate and an N-linked glycan that monomeric UGGT can re-glucosylate on the same glycoprotein. MD simulations estimate the Parodi limit to be around 60-70 Å. Re-glucosylation assays using UGGT deletion mutants suggest that the TRXL2 domain is necessary for activity against urea-misfolded bovine thyroglobulin. Taken together, our findings support a “one-size-fits-all adjustable spanner” substrate recognition model, with a crucial role for the TRXL2 domain in the recruitment of misfolded substrates to the enzyme’s active site.

## Introduction

A wonderfully efficient protein folding machinery in the Endoplasmic Reticulum (ER) of eukaryotic cells ensures that only correctly folded glycoproteins can exit the ER, proceed to the Golgi, and from there continue along the secretory pathway towards their cellular or extracellular destinations [1]. The stringency of this Endoplasmic Reticulum Quality Control (ERQC) system is of great advantage to healthy cells because it allows time for complex glycoproteins to fold in the ER and prevents premature secretion of incompletely folded species. In the background of a misfold-inducing missense mutation in a secreted glycoprotein gene, the resulting misfolded glycoprotein is either retained in the ER by ERQC or degraded by the ER associated degradation (ERAD) machinery [2]. ERQC bears particularly unfortunate consequences when the mutation induces a minor folding defect but does not abrogate the function of the glycoprotein (“responsive mutant”): in this case ERQC causes disease by blocking the secretion of the glycoprotein mutant, even though its residual activity would be beneficial to the organism (see for example [3]).

Central to ERQC is the ER-resident 170 kDa enzyme UDP-Glucose:glycoprotein Glucosyl Transferase (UGGT). The enzyme selectively re-glucosylates a misfolded glycoprotein on one of its N-glycans and promotes its association with the ER lectins calnexin and calreticulin, thus mediating ER retention. Only correctly folded glycoproteins escape UGGT-mediated re-glucosylation and can progress down the secretory pathway – to the Golgi and beyond. More than 25 years after the discovery of UGGT [3], recent structural and functional work has uncovered the protein’s multi-domain architecture and obtained preliminary evidence of its inter-domain conformational flexibility [4–6]. Negative-stain electron microscopy and small-angle X-ray scattering (SAXS) first revealed an arc-like structure with some degree of structural variability [4]. Soon after, four distinct full length *Chaetomium thermophilum* UGGT (*Ct*UGGT) crystal structures, together with a 15 Å cryo-EM reconstruction of the same protein, suggested that the C-terminal portion - comprising two β-sandwiches and the catalytic domain - constitutes a relatively rigid structure, while most of the conformational flexibility localizes to the four N-terminal thioredoxin-like (TRXL) domains [5]. Contrary to these observations, relative flexibility of the catalytic domain with respect to the rest of the structure was proposed by a different study of *Thermomyces dupontii* UGGT (*Td*UGGT), based on atomic force microscopy data and a 25 Å negative stain EM map, to which crystal structures for the catalytic domain and the remaining part of *Td*UGGT were separately fitted [6, 7].

Here, we further characterize UGGT’s inter-domain flexibility and seek to clarify the above-mentioned controversy regarding the relative movements of the C-terminal vs. N-terminal portions of the molecule. The question is important in order to understand the molecular basis for the enzyme’s promiscuity. We analyze the available negative stain EM and crystal structures, carry out Molecular Dynamics (MD) experiments and describe four new *Ct*UGGT crystal structures. We find that the majority of UGGT’s overall inter-domain motions can be depicted in terms of three types of main movements of the N-terminal misfold recognition domains, which we call bending, twisting and clamping. Our results are consistent with the catalytic domain being tightly associated with the β-sandwiches, thus ruling out major relative flexibility of the C-terminal domain with respect to the N-terminal ones. The second TRXL domain (TRXL2) proves to be the most mobile one. Kinetic measurements on TRXL2 and TRXL3 deletion UGGT mutants in re-glucosylating assays of urea-misfolded bovine thryoglobulin suggest that only the former domain is absolutely necessary for re-glucosylation of this substrate. We discuss the functional implications of these discoveries.

## Results

### A new crystal structure of *Ct*UGGT adds to the landscape sampled by previously observed UGGT conformations

In the previously reported crystal *Ct*UGGT structures, a full length eukaryotic UGGT revealed four thioredoxin-like domains (TRXL1-4) arranged in a long arc, terminating in two β–sandwiches (βS1 and βS2) tightly clasping the glucosyl-transferase family 24 (GT24) domain [5]. The wild-type protein was captured in three different conformations, called ‘open’ (PDB ID 5MZO), ‘intermediate’ (PDB ID 5MU1) and ‘closed’ (PDB ID 5N2J). Additionally, the double Cysteine mutant D611C-G1050C, engineered to form an extra disulfide bridge between the TRXL2 and βS2 domains, was trapped in a ‘closed-like’ conformation (PDB ID 5NV4). Those four *Ct*UGGT structures mainly differ in the spatial organization of domains TRXL2 and TRXL3 (Figure 1, and SI Appendix movie I). Across these structures, the TRXL2 domain is rotated by different amounts with respect to the rest of the protein and adopts different degrees of proximity to it. The TRXL3 domain instead appears in the same relative conformation in all structures, except for the ‘open’ one (right-hand side panel in Figure 1A and green in Figures 1B,C), in which the TRXL3 and TRXL1 domains move apart, leading to the opening of a cleft between them.

**Figure 1.**
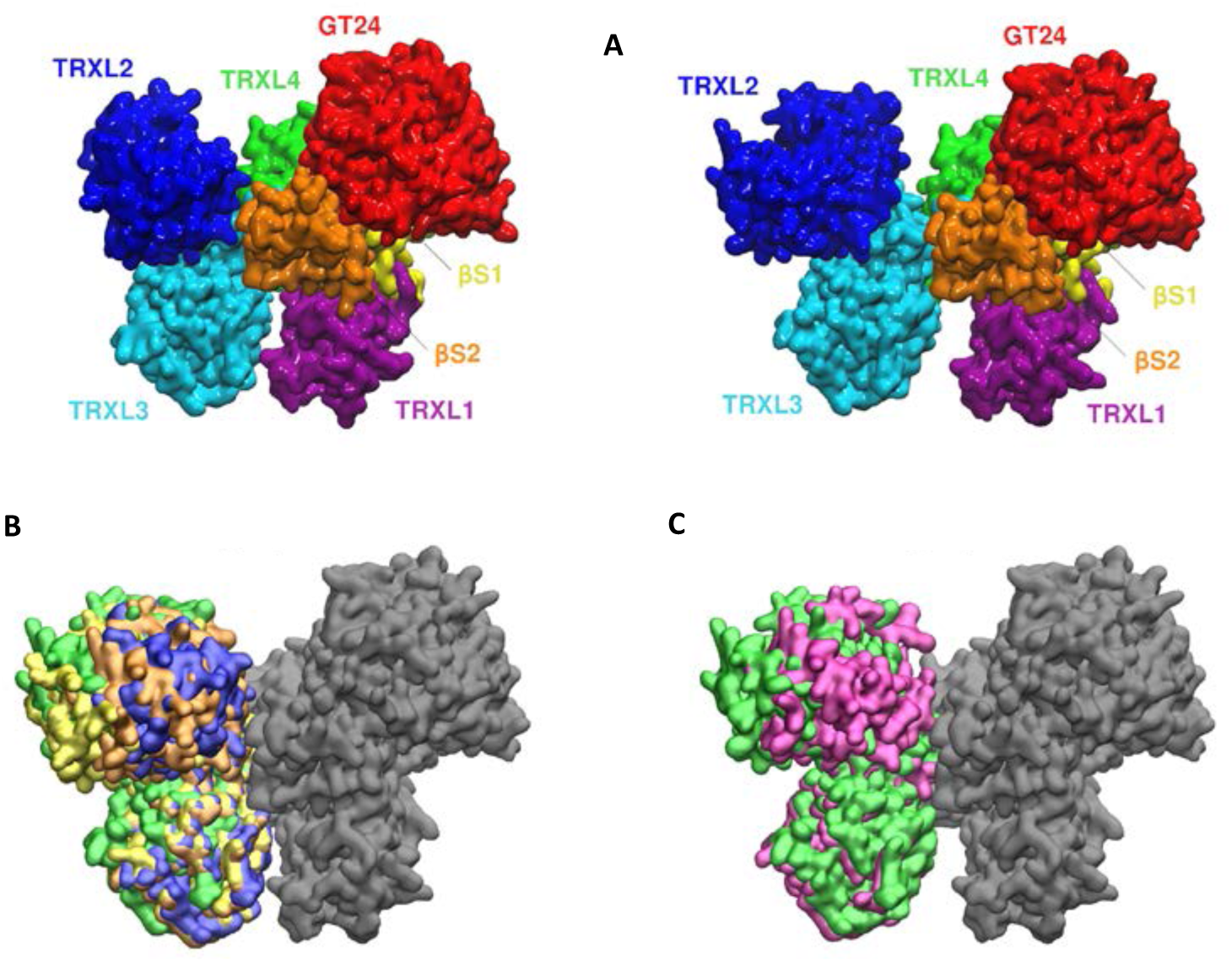
A new crystal structure of WT *Ct*UGGT bears similarity to previously observed closed and open conformations. **(A)** Structural comparison between *Ct*UGGT in “closed” (PDB ID: 5N2J, left hand side) and “open” (PDB ID: 5MZO, right hand side) conformations, coloured domain by domain: TRXL1 (residues 45-220): magenta; TRXL2 (residues 414-656): blue; TRXL3 (residues 667-880): cyan; TRXL4 (residues 275-410; 897-950): green; βS1 (residues 28-36; 225-242; 957-1037): yellow; βS2 (residues 1039-1149): orange; GT24 (residues 1197-1475): red. **(B)** Superimposition of all four *Ct*UGGT X-ray structures available prior to this publication; domains coloured in grey (GT24, βS1, βS2 and TRXL4) represent the relatively rigid portion of the molecule (RMSD_C_α less than 0.750 Å) which was used to align the structures. TRXL2 and TRXL3 domains are coloured as follows: blue, ‘closed’ conformation (PDB ID: 5N2J); orange, D611C-G1050C mutant *aka* ‘closed-like’ conformation (PDB ID: 5NV4); yellow, ‘intermediate’ conformation (PDB ID: 5MU1); green, ‘open’ conformation (PDB ID: 5MZO). **(C)** Superimposition of the ‘open’ conformation (TRXL2 and TRXL3 domains in green) with the recently reported ‘new intermediate’ *Ct*UGGT_Kif_ conformation (TRXL2 and TRXL3 domains in magenta) (PDB ID: 6TRF).

We describe here a fifth novel full-length *Ct*UGGT structure (hereinafter *Ct*UGGT_Kif_, PDB ID 6TRF), obtained from cells treated with the mannosidase inhibitor kifunensine (the compound prevents elaboration of N-linked glycans along the secretory pathway and ensures that secreted glycoproteins carry high-mannose glycans). *Ct*UGGT_Kif_ adopts a conformation which combines a TRXL1-TRXL3 distance as found in the ‘open’ conformation, but a TRXL2/TRXL3 relative orientation similar to the one found in the ‘close j ~÷|”Ji u d-like’ conformation (Figure 1C). We label this *Ct*UGGT_Kif_ conformation ‘new intermediate’.

In order to establish a framework for the discussion of the motions of the UGGT molecule, mapping its inter-domain conformational landscape in a compact way, we define here 3 collective conformational coordinates (CCs). ‘CC1’ or ‘clamp’, describes the changes in the distance between the centres of mass of the TRXL1 and TRXL3 domains, and measures the openness of the cleft between them; ‘CC2’ or ‘bend’, describes the changes in the angle between the centres of mass of the TRXL1, TRXL2 and TRXL3 domains, and measures the proximity of the TRXL2 and GT24 domains across the central saddle; lastly, ‘CC3’ or ‘twist’, describes the changes in the dihedral angle between the Cα atoms of residues Y518, F466, T863 and I735 (the first two residues in the TRXL2 and the last two in the TRXL3 domain), thus informing on the relative orientation of the TRXL2 and TRXL3 domains.

Figure 2 reports the values of the conformational coordinates for the conformations observed in *Ct*UGGT X-ray structures. The pair of CC1/clamp and CC3/twist values for the ‘new intermediate’ *Ct*UGGT_Kif_ structure, compared with the ones for the previous structures, suggests that clamping and twisting motions may be to an extent independent, and the molecule exist in states where the TRXL1:TRXL3 domain clamp is open, *Ct*UGGT_Kif_ (PDB ID 6TRF) CC1 =43.2 Å, while at the same time preserving a middle-of-the-range value for the TRXL2:TRXL3 domain pair twist: *Ct*UGGT_Kif_ (PDB ID 6TRF) CC3=3.2°.

**Figure 2.**
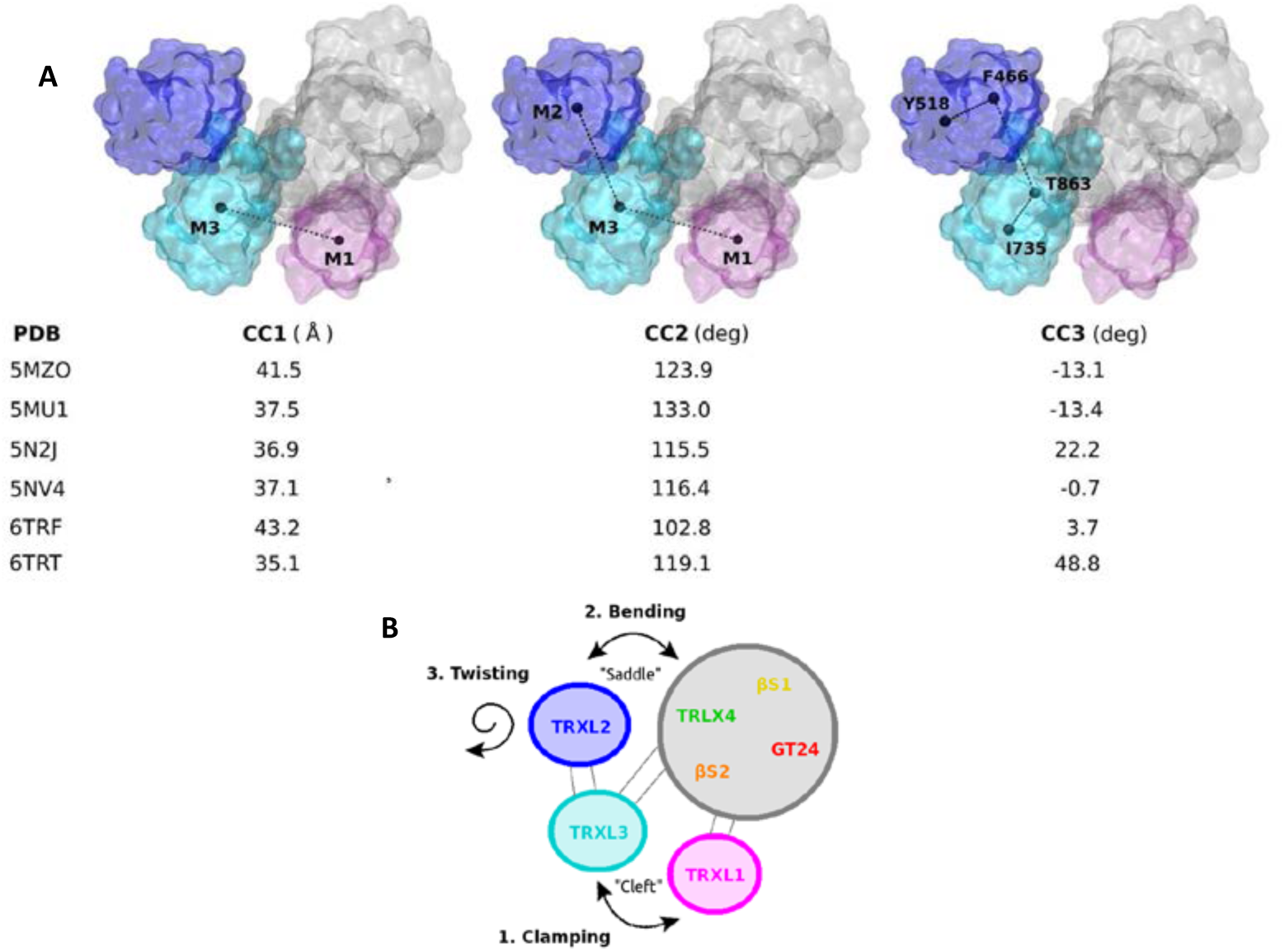
Three main UGGT motions. **(A)** Conformational coordinates (CCs) for describing the *Ct*UGGT conformational states observed in the X-ray structures; domain colour code is the same as in figure 1: TRXL1 (magenta), TRXL2 (blue) and TRXL3 (cyan). Along ‘CC1’, the ‘clamp’ coordinates measures the openness of the cleft between the TRXL1 and TRXL3 domains; along ‘CC2’, the ‘bend’ coordinate measures the distance between the TRXL2 and GT24 domains across the central saddle; along ‘CC3’ the ‘twist’ coordinate changes with the relative orientation of the TRXL2 and TRXL3 domains. **(B)** Simplified representation of *Ct*UGGT overall movements. ‘Clamping’ movement between domains TRXL3 and TRXL1; bending’ movement between TRXL2 and the core comprised of domains ‘GT24-βS1-βS2-TRXL4’; ‘twisting’ movement of TRXL2 with respect to TRXL3. The grey area represents the strong structural inter-domain orientation invariance of the TRXL4-βS1-βS2-GT24 domains.

### UGGT’s motions can be described in simple terms as two rigid groups of domains moving with respect to each other

We asked the question whether the conformational landscape spanned by UGGT full-length crystal structures can be extended by *in silico* molecular dynamics. We performed 250 ns long MD simulations starting from four of the *Ct*UGGT crystal structures and computed principal components (PCs, also called essential modes [8]) from the four individual MD trajectories and from the fusion of all four MDs into a single trajectory. UGGT MD trajectories, as expected, span a wider conformational landscape compared to the set of crystal structures. Overall, UGGT’s motions can be described in simple terms as two rigid groups of domains moving with respect to each other: one group is formed by domains TRXL2-TRXL3; and the other is formed by domains TRXL1-TRXL4-βS1-βS2-GT24 – the latter group is enclosed in a grey circle in Figure 2B. The interface between domains TRXL3 and TRXL4 acts as a hinge region between the two domain groups.

The first two PCs of the joint MD simulation suffice to parameterize most of the observed motions: PC1 (Figure 3A, left hand side panel) describes the transition between ‘open’ and ‘closed’ states and follows domain TRXL2 bending towards domain βS2 across the central saddle, with TRXL3 and TRXL1 clamping together across the cleft at the same time (SI Appendix Figure S1 A and SI Appendix Movie II). Figures 3B and 3C show that the MD simulations starting from the ‘open’ and ‘intermediate’ crystal structures both move significantly along PC1 and visit both ‘open’ and ‘closed’ states. The MD simulation starting from the ‘open’ structure shows a back-and-forth movement along PC1 (Figure 3C), while the one starting from the ‘intermediate’ structure drifts to the ‘closed’ state and beyond (Figure 3B).

**Figure 3.**
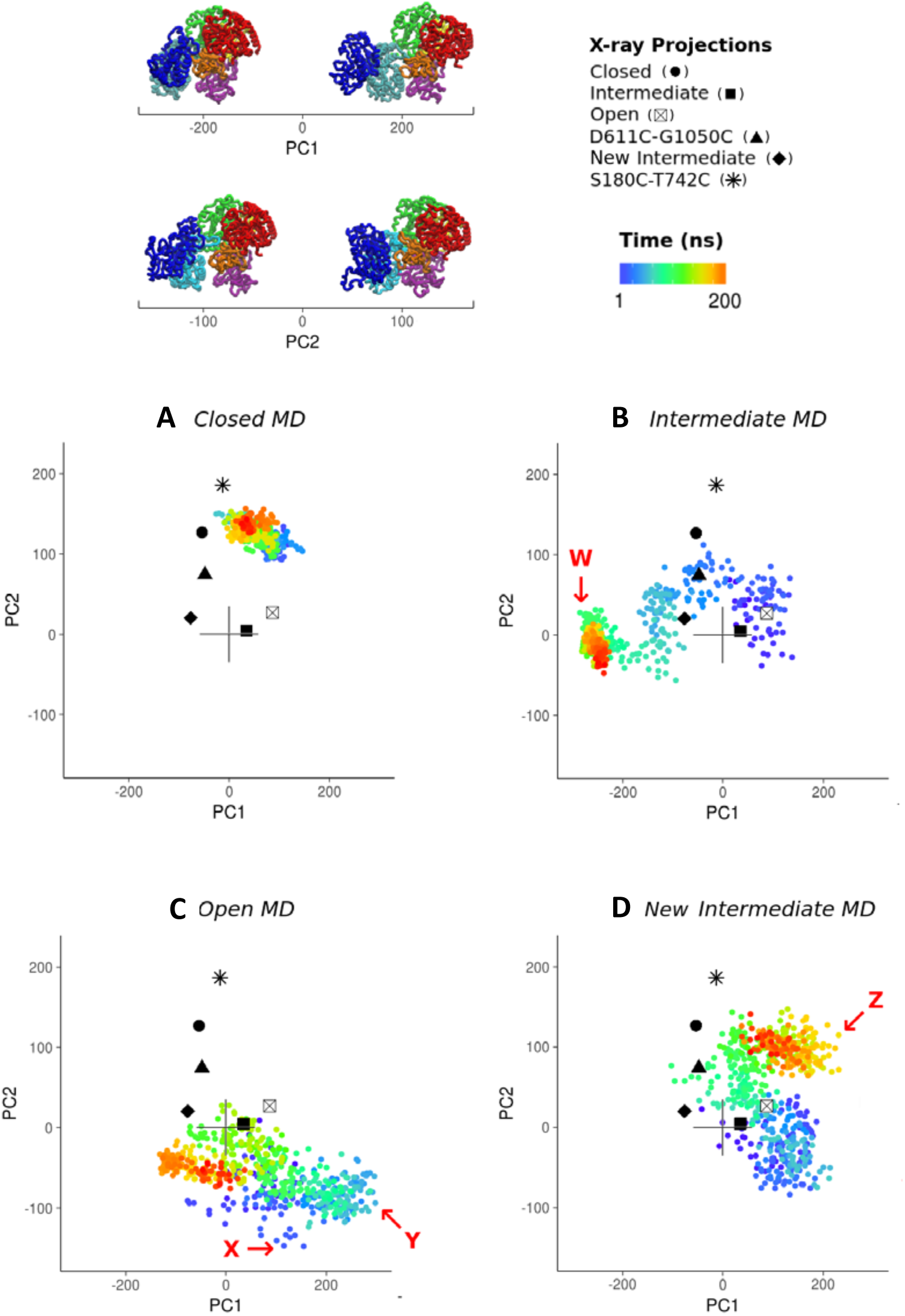
Relative movements of the TRXL1-TRXL4-βS1-βS2-GT24 domains with respect to the TRXL2-TRXL3 domains. Projections of individual MD trajectories and their respective X-ray starting structures onto the full conformational landscape as described by the first and second PCs, coloured as a function of time. Domains coloured as in Figure 1A. IN red we list a few *Ct*UGGT structures representative of extreme values of the conformational coordinates (CCs), as identified within the MD conformational landscape (see also Figure 5).

PC2 (Figure 3A, right hand side panel) describes a movement in which the TRXL2 domain rotates with respect to TRXL3 (SI Appendix Figure S1 B and SI Appendix Movie III), and the βS2, TRXL1 and TRXL4 domains also undergo motion. The motion encoded by PC2 is well represented in the MD starting from the ‘new intermediate’ *Ct*UGGT_Kif_ structure, whose projection in Figure 3D also shows a considerable degree of back-and-forth movement.

The fact that the *Ct*UGGT βS1-βS2:GT24 portion of the molecule behaves as one relatively rigid structure throughout the MD simulations is hardly surprising: the domains bury a 1400 Å^2^ surface, with a calculated −7.1 kcal/mol solvation free energy gain [9]. The βS1-βS2:GT24 interface is supported by 16 hydrogen bonds, five salt bridges, and 11 hydrophobic interactions, involving 86 residues overall. The PISA server Complex Formation Significance Score (CSS) is 1.0 [9], suggesting that the contacts in the *Ct*UGGT βS1-βS2:GT24 interface are sufficient to support the observed N_term_:C_term_ interdomain structure. The solvation free energy gain computed by the same server has a P-value of 0.326 (P<0.5 indicates interfaces with “surprising” - *i.e.* higher than average - hydrophobicity, implying that the interface is likely interaction-specific) [9].

The tight association we observe between the GT24 and βS1-βS2 domains is at odds with a hypothesis formulated in 2017 and based on negative stain EM and atomic force microscopy (AFM) of *Thermomyces dupontii* UGGT (*Td*UGGT): the study proposed that the UGGT GT24 domain assumes a number of different relative orientations with respect to the rest of the molecule, enabled by the flexible linker between the βS2 and GT24 domains [6, 7]. Of the 48 residues contributing side chains to the UGGT βS1-βS2:GT24 interface, 44 are conserved between *Td*UGGT and *Ct*UGGT, and none of the 4 side chain differences would likely abrogate contributions to the interdomains interface (see SI Appendix Figure S4). This prompted us to hypothesize that the GT24 and βS1-βS2 domains constitute a rigid group in *Td*UGGT also (and, by extension, in UGGTs across all eukaryotes), just as observed in full-length *Ct*UGGT structures and their MD simulations. In absence of a full-length *Td*UGGT crystal structure, the only information about the relative orientation of *Td*UGGT GT24 and βS1-βS2 domains comes from a 25 Å negative stain EM reconstruction of *Td*UGGT in complex with an anti-*Td*UGGT antibody fragment (Fab) [6, 7]. In order to check if the *Td*UGGT negative stain EM reconstruction is compatible with a model in which GT24 and βS1-βS2 domains also form a rigid group, we generated a full-length *Td*UGGT homology model, selected a representative Fab structure from the Protein Databank, and proceeded to fit them into the 25 Å negative-stained EM map for the complex of *Td*UGGT with its Fab (separately fitting the models into both the original map and its enantiomeric mirror image [10]). The correlation coefficients (CCs) between the 25 Å negative-stained EM map and the *Td*UGGT and Fab models are around 90% for both original hand (Figure 4A-C) and inverted hand map (Figure 4D-F), for both *Td*UGGT and Fab models. In the fitted models, the Fab contacts the 440-460 portion of *Td*UGGT domain TRXL2, in agreement with the published Fab epitope (residues *Td*UGGT 29-468) [6, 7]. Therefore, the 25 Å negative-stained EM map of the complex of *Td*UGGT with its Fab can be fitted by a full-length *Td*UGGT model without invoking any detachment of the catalytic domain from the βS1-βS2 region, contrary to what stated in [6, 7].

**Figure 4.**
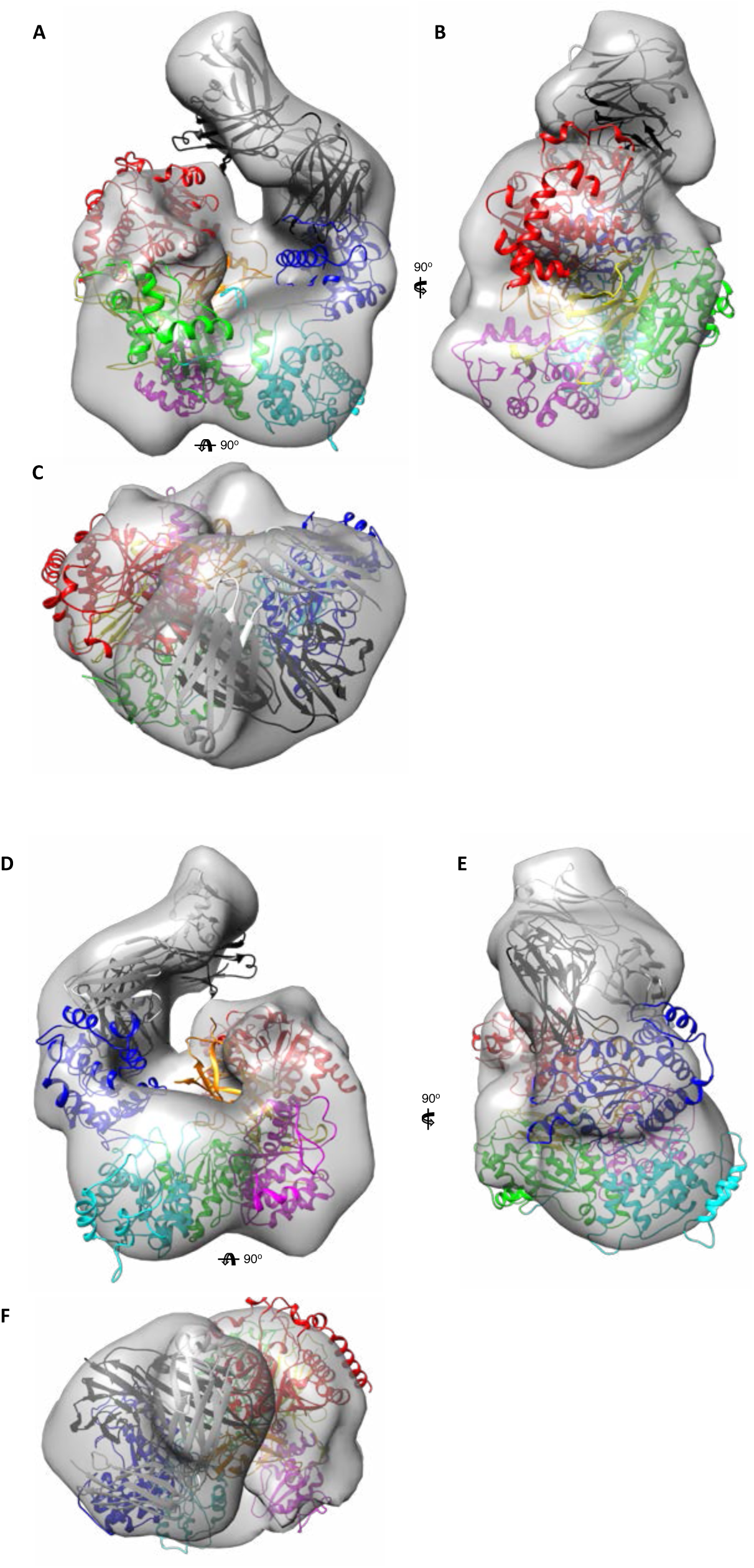
Fitting of full-length *Td*UGGT and Fab models in the negative-stain EM map for the complex of *Td*UGGT and an anti-*Td*UGGT Fab. The homology model for *Td*UGGT (residues 29-1466) was coloured as follows: TRXL1 (residues 40-219): magenta; TRXL2 (residues 413-657): blue; TRXL3 (residues 658-898): cyan; TRXL4 (residues 239-412; 899-958): green; βS1 (residues 29-39; 220-238; 959-1037): yellow; βS2 (residues 1038-1151): orange; GT24 (residues 1190-1466): red. A generic Fab structure was chosen for the fitting of the anti-*Td*UGGT Fab antibody fragment (PDB ID 1FGN, 214+214 residues, MW=46927 Dalton), painted black (heavy chain) and white (light chain). The 25 Å negative-stain EM map is contoured at a contour level appropriate for enclosing the mass of the *Td*UGGT plus a Fab fragment (i.e. about 356,800 Å^3^ corresponding to a mass of 295,000 Dalton, based on a specific volume of 1.21 Å per Dalton [51]). A, B and C: three views of the *Td*UGGT and Fab models fitted in the original hand of the negative stain EM map (with B and C rotated by 90’ with respect to view A around the centre of mass of the model, along the vertical and horizontal direction, respectively). D, E and F: three views of the same *Td*UGGT and Fab models fitted to the inverse hand of the negative stain EM map (with E and F rotated by 90’ with respect to view D around the centre of mass of the model, along the vertical and horizontal direction, respectively).

### UGGT ‘Twisting’ and ‘clamping’ motions are uncorrelated

As shown in Figure 3, MD simulations take *Ct*UGGT beyond the space sampled by the X-ray structures. In particular, the MD simulations starting from the ‘open’, ‘intermediate’ and ‘new intermediate’ *Ct*UGGT_Kif_ structures reach conformations with extreme PC values (Figure 5). Most notably, the structure labelled ‘W’ (Figure 5A) represents an extreme version of a closed state. It reveals that the 7 UGGT domains can converge to a conformation of very compact overall shape. At the opposite end of the UGGT conformational landscape, structures labelled ‘X’, ‘Y’ and ‘Z’ (Figures 5B-D) resemble open-like states. Structure ‘X’ in particular (Figure 5B) presents a notable opening of the TRXL1-TRXL3 cleft along the clamping motion described by CC1, while also showing a considerable degree of twisting along CC3. In contrast to ‘X’, structures ‘Y’ and ‘Z’ (Figures 5C,D) both exhibit a clamped cleft, but at extreme CC2 values, suggesting that UGGT is able to push the bending motion even further than observed in the ‘open’ structure, while at the same time retaining a clamped cleft. All together, the set of CtUGGT crystal structures and their MD simulations confirm that ‘twisting’ and ‘clamping’ UGGT motions are uncorrelated, as we first hypothesized upon determination of the *Ct*UGGT_Kif_ structure.

**Figure 5.**
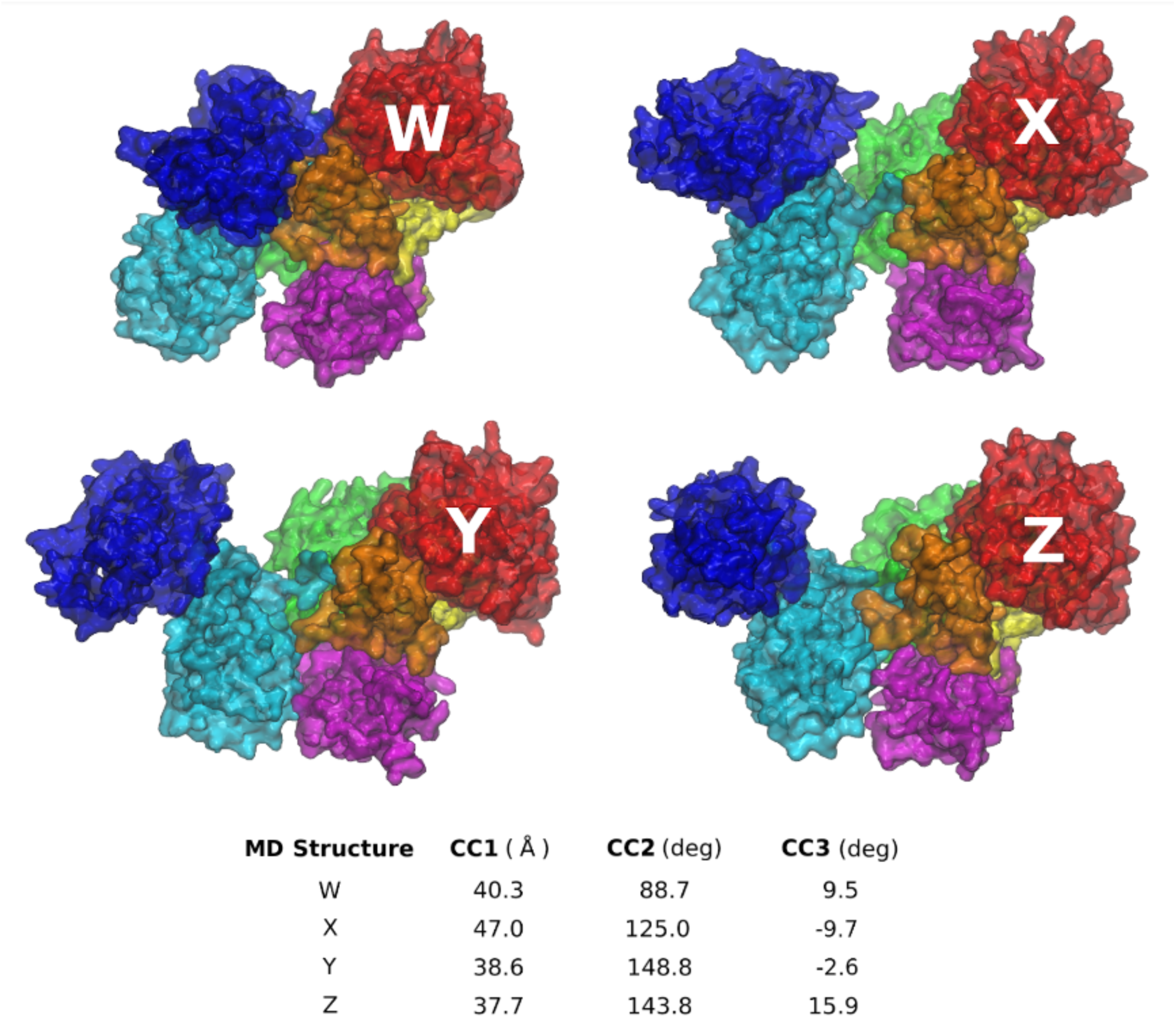
MD snapshots with extreme PC values, and their respective Conformational Coordinates measured. A few *Ct*UGGT structures representative of extreme values of the conformational coordinates (CCs), as identified within the MD conformational landscape. ‘W’: ‘clamped, bent and twisted shut’ (small values of CC1, CC2 and CC3). ‘X’: ‘clamped open’ and ‘twisted open’ (large CC1 and CC3 values); ‘Y’ and ‘Z’: ‘clamped shut’ (smaller values of CC1) but ‘bent open’ (large CC2 values). Domain coloured as in Figure 1A.

### UGGT activity is underpinned by its inter-domain conformational mobility

The study in [5] engineered two *Ct*UGGT double cysteine mutants, *Ct*UGGT^N796C/G1118C^ and *Ct*UGGT^D611C/G1050C^, designed to form disulfide bridges across the interfaces of the TRXL3-βS2 and TRXL2-βS2 domains, respectively. Due to the extra disulfide bridge, *Ct*UGGT^N796C/G1118C^ cannot attain the ‘open’ state, while *Ct*UGGT^D611C/G1050C^ cannot attain either the ‘open’ or the ‘intermediate’ conformation. As evidenced in the SI Appendix Figure S1A, the MD trajectory starting from the *Ct*UGGT^D611C/G1050C^ structure shows significantly restricted mobility along the first PC, confirming that the extra disulphide bridge in *Ct*UGGT^D611C/G1050C^ tethers the TRXL2 and βS2 domains in a closed conformation; along the second PC, *Ct*UGGT^D611C/G1050C^ moves further than the other structures. The *Ct*UGGT^N796C/G1118C^ mutant on the other hand still retains most of its mobility, being able to explore a similar conformational space as those observed for wild-type *Ct*UGGT (SI Appendix Figure S1B).

In activity assays of UGGT-mediated re-glucosylation of the misfolded glycoprotein substrate urea-misfolded bovine thyroglobulin, both *Ct*UGGT^N796C/G1118C^ and *Ct*UGGT^D611C/G1050C^ mutants had lower activity than wild-type *Ct*UGGT, but *Ct*UGGT^N796C/G1118C^ had a higher catalytic activity and a lower melting temperature than *Ct*UGGT^D611C/G1050C^ [5]. Taken together, these results suggested that the ‘bending’ motion is important for re-glucosylation of this particular substrate. In order to probe the functional role of the ‘clamping’ motion uncovered in the present analysis, we engineered three novel double cysteine *Ct*UGGT mutants: *Ct*UGGT^G177C/A786C^, *Ct*UGGT^G179C/T742C^ and *Ct*UGGT^S180C/T742C^, all designed to form disulfide bridges across the TRXL1 and TRXL3 domains, clamping the cleft between them shut. All three *Ct*UGGT double Cys mutants were expressed and purified from the supernatant of mammalian HEK293F cells; the presence of the engineered disulfide bridges was confirmed by mass spectrometry (SI Appendix Figure S3). The crystal structures of *Ct*UGGT^G177C/A786C^ and *Ct*UGGT^S180C/T742C^ were determined to about 4.7-4.5 Å resolution. Both crystal structures show the TRXL3 domain tethered to the TRXL1 domain by the extra disulfide bridge (Figure 6A). We tested the *in vitro* activity of the three double Cys mutants (in addition to the activity of the WT and the already published *Ct*UGGT^D611C/G1050C^) in a re-glucosylation assay of the UGGT substrate urea-misfolded bovine thyroglobulin (Figure 6B). Despite their structural similarity, the *Ct*UGGT^S180C/T742C^ and *Ct*UGGT^G177C/A786C^ mutants differ significantly in their ability to re-glucosylate urea-misfolded bovine thyroglobulin: the former is more active than WT *Ct*UGGT, while the latter has similar activity to it.

**Figure 6.**
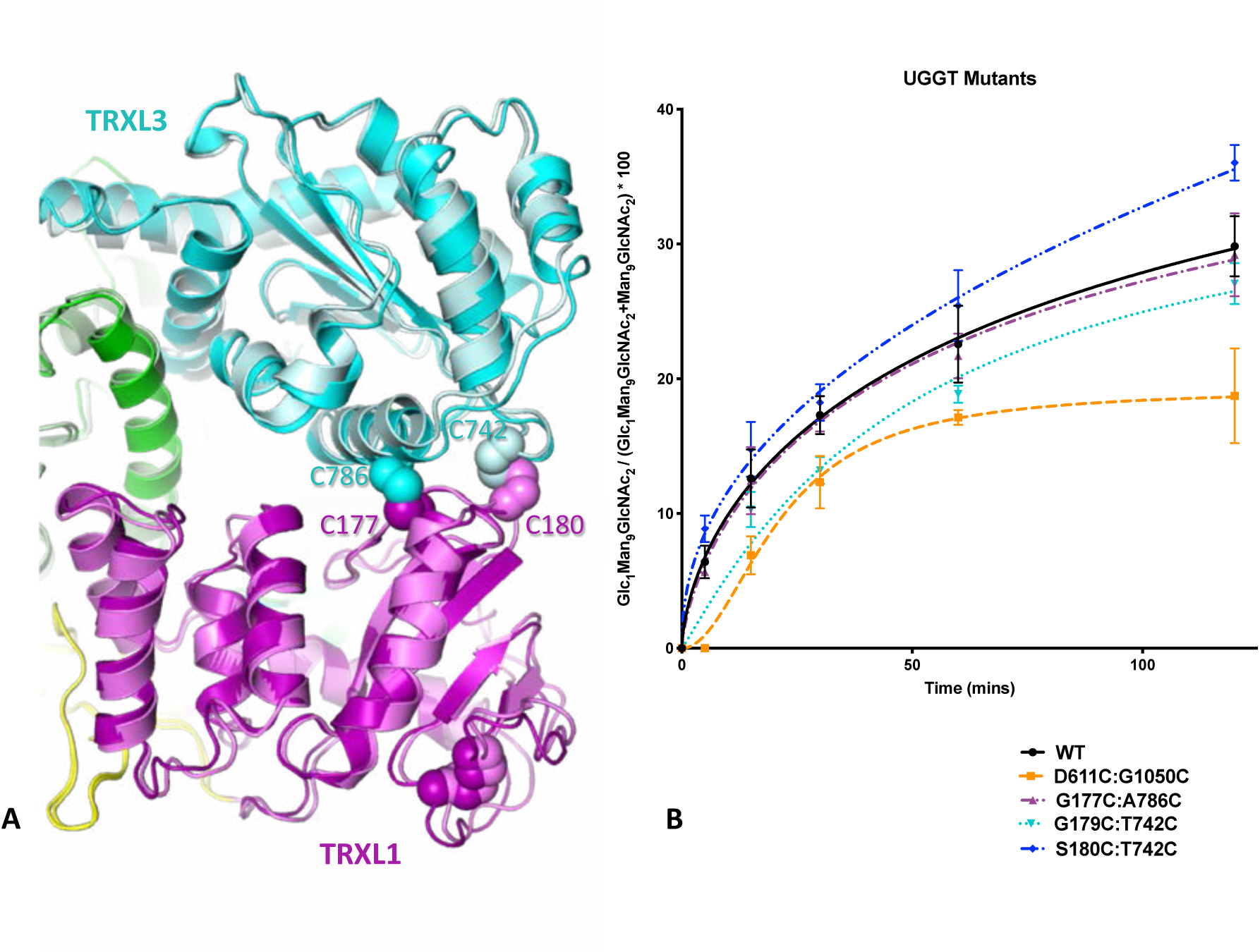
Activity of *Ct*UGGT double Cys mutants. **(A)** The TRXL1 (magenta) and TRXL3 (cyan) domains in the crystal structures of *Ct*UGGT^G177C/A786C^ (PDB ID: XXXX, dark colours) and *Ct*UGGT^S180C/T742C^ (PDB ID: 6TRT, lighter colours). The disulfide bonds are in spheres representation. **(B)** Re-glucosylating activity of *Ct*UGGT double Cys mutants and WT *Ct*UGGT against urea-misfolded bovine thyroglobulin.

### *Ct*UGGT-mediated re-glucosylation of urea-misfolded bovine thyroglobulin requires the TRXL2 domain

In order to probe the contributions of individual UGGT TRXL domains to UGGT re-glucosylating activity, we cloned three mutants of *Ct*UGGT, each lacking one of the TRXL1-3 domains: *Ct*UGGT-ΔTRXL1, lacking residues 42-224; *Ct*UGGT-ΔTRXL2, lacking residues 417-650; and *Ct*UGGT-ΔTRXL3, lacking residues 666-870. Only the latter two mutants expressed and were purified, and *Ct*UGGT-ΔTRXL2 was the only TRXL domain deletion mutant that yielded crystals, enabling crystal structure determination by X-ray diffraction to 5.7 Å resolution. The *Ct*UGGT-ΔTRXL2 crystal structure most closely resembles the ‘closed’ structure (1.32 Å rmsd_Cα_ with PDB ID 5NV4, over 975 residues) apart from a minor rearrangement of the TRXL3 domain, which moves away from the rest of the truncated molecule (Figure 7A). UGGT-mediated re-glucosylation activity assays of *Ct*UGGT-ΔTRXL2 and *Ct*UGGT-ΔTRXL3 against urea-misfolded bovine thyroglobulin detect impaired re-glucosylation activity upon deletion of TRXL3 and complete loss of activity upon deletion of TRXL2 (Figure 7B).

**Figure 7.**
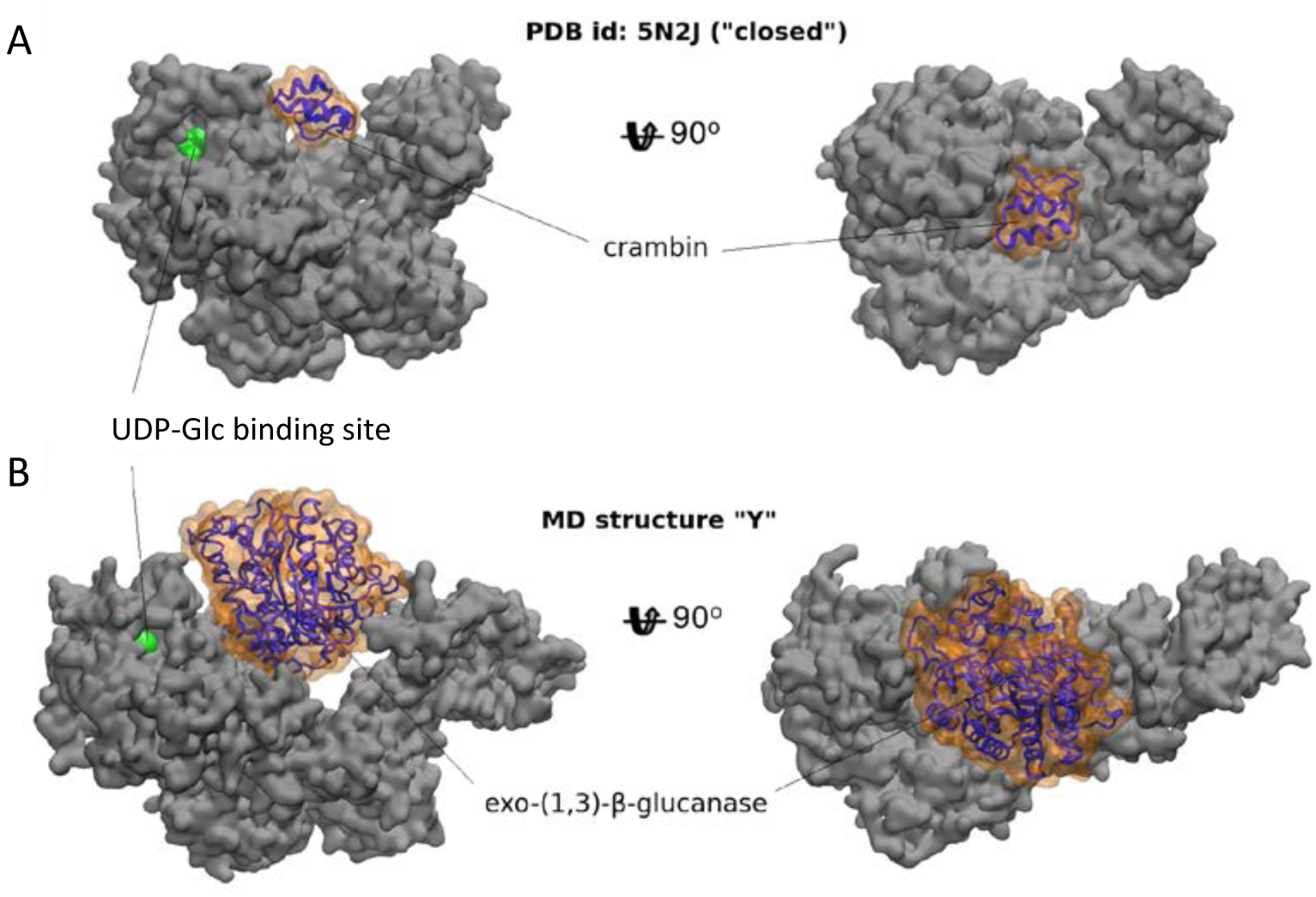
*Ct*UGGT ΔTRXL2 and ΔTRXL3 deletion mutants. **(A)** Crystal structure of *Ct*UGGT-ΔTRXL2 (PDB ID: 6TS2, copy “A”, solid colours) overlayed onto WT *Ct*UGGT (‘open’ conformation (PDB ID: 5MZO), semi-transparent). Domain coloured as in Figure 1A. **(B)** Activity of *Ct*UGGT-ΔTRXL2 and *Ct*UGGT-ΔTRXL3 against urea-misfolded bovine thyroglobulin, compared to WT *Ct*UGGT.

### UGGT: a ‘one size fits all’ adjustable spanner?

Taken together, our UGGT dynamics, structures and functional data so far suggest an “adjustable spanner” model in which the enzyme’s central saddle may bind the misfolded portion of substrate glycoproteins, with the TRXL2 domain providing most of the conformational variability, and various UGGT conformations adapting to different sizes/shapes of glycoprotein substrates [4, 5]. Measurements of the central saddle surface area (SA) in the observed UGGT MD conformations, from the most compact structure, ‘W’, to most open structure, ‘Y’, span the range 8,600-11,300 Å^2^, with average values around 9,200-9,700 Å^2^ for most crystal structures (SI Appendix movie IV). Substrate glycoproteins with a ‘radius of gyration’ (rog) ⪟ 15 Å and around 150-200 residues or less, would snugly fit in the central saddle of compact or middle-of-the-range UGGT conformers (Table 1 and Figure 8A). In contrast, for binding of larger substrates (15 Å ⪟ rog ⪟ 23 Å, and 200-500 residues) an opening of the central saddle would be needed (Figure 8B).

**Figure 8.**
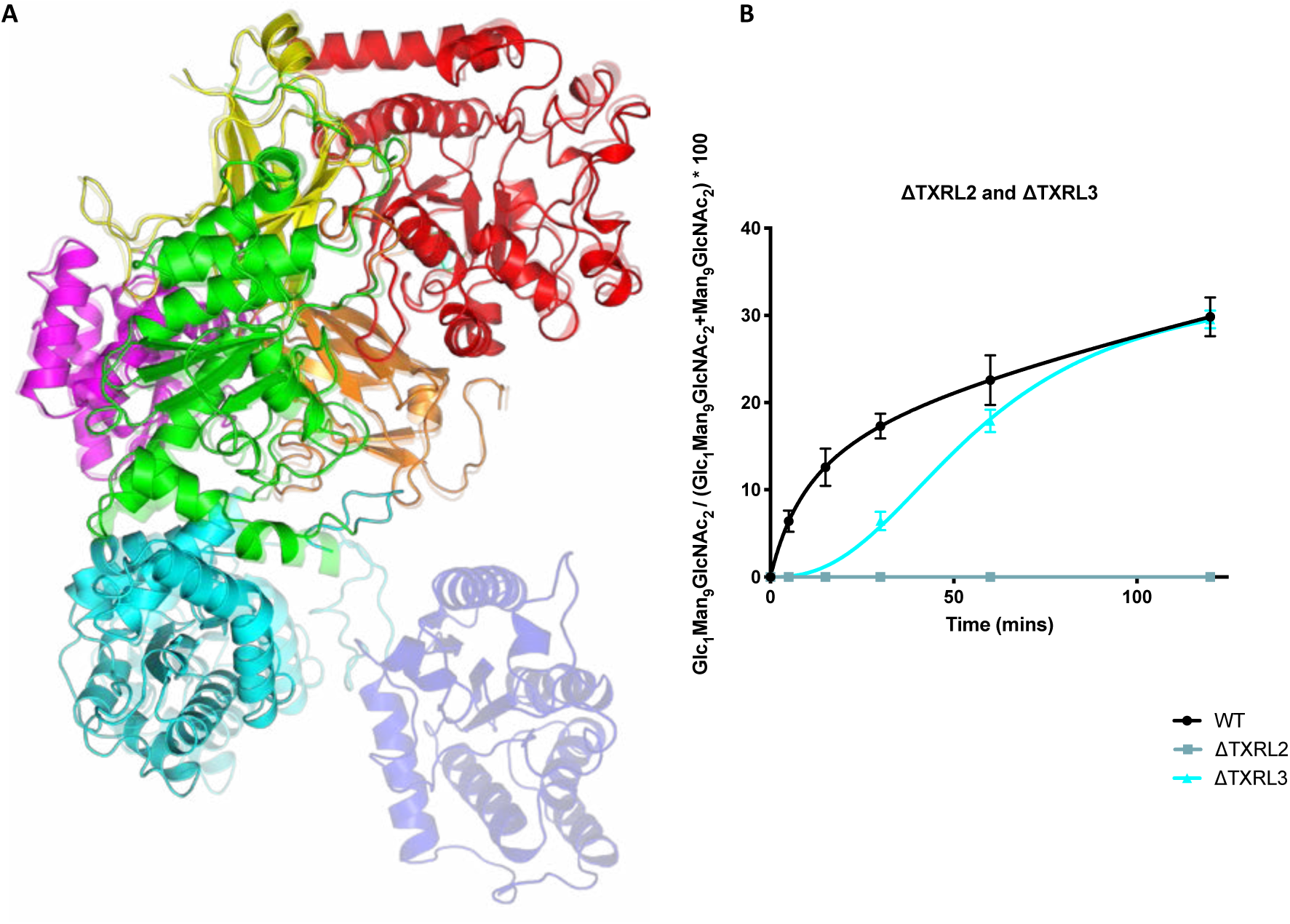
The UGGT ‘one-size-fits-all’ adjustable spanner model. Two *Ct*UGGT conformations in complex with experimentally-validated substrates of different sizes. The bright green region shows the active site. **(A)** Crambin in complex with *Ct*UGGT “closed” crystal structure, conformation, MD-derived structure *Ct*UGGT ‘W’ of Figure 5. **(B)** Exo-(1,3)-β-glucanase in complex with MD-derived structure *Ct*UGGT ‘Y’ of Figure 5.

**Table 1.** In vitro UGGT substrates. List of various UGGT misfolded glycoprotein substrates described in the literature as UGGT substrates in *in vitro* experiments. We have not included glycoproteins that have been inferred to be UGGT substrates by *in cellula* experiments (see for example [25, 61–64]) nor glycoproteins that are *bona fide* UGGT substrates but whose structure has not been determined [11, 65]). (*): structures are available for the mature glycoprotein but it is the pro-glycoprotein (previous to protease cleavage) that folds in the ER under UGGT control – so we have not estimated the RoG.

| Substrate | PDB ID | Number of residues | Radius of Gyration (Å) | Reference |
| --- | --- | --- | --- | --- |
| *Crambe hispanica* crambin | 1CRN | 46 | 9.7 | [54] |
| *Hordeum vulgare* chymotrypsin inhibitor 2 | 2CI2 | 64 | 11.4 | [29] |
| Human interleukin 8 (IL-8) | 1ICW | 72 | 12.5 | [55] |
| Human prosaposin A | 5NXB | 87 | N/A<sup>(*)</sup> | [56] |
| Bovine RNase BS-prot | 2E33 | 104 | 14.7 | [14, 15] |
| Bovine RNase B | 2E33 | 124 | 14.3 | [14, 15, 57] |
| *Staphylococcus aureus* nuclease | 1NUC | 149 | 14.4 | [14, 15, 57] |
| *Trypanosoma cruzi* cruzipain | 3I06 | 215 | N/A<sup>(*)</sup> | [58] |
| Soybean agglutinin | 4D69 | 234 | 16.9 | [23] |
| Human alpha-galactosidase | 3HG5 | 390 | 21.5 | [59] |
| Human exo-(1,3)-β-glucanase | 1H4P | 407 | 20.3 | [13] |
| Human transferrin | 6D04 | 678 | 29.7 | [60] |

## Discussion

Since the discovery of UGGT back in 1989 [11, 12], activity studies have used a range of glycoprotein substrates, such as urea-misfolded bovine thyroglobulin [11], mutants of exo-(1,3)-β-glucanase [13], RNase BS [14, 15], small size synthetic compounds bearing high-mannose glycans attached to fluorescent aglycon moieties such as ‘TAMRA’ and ‘BODIPY’ [16, 17] and chemically synthesized misfolded glycoproteins [18–20], to mention only a few. Although a comprehensive list of physiological UGGT substrate glycoproteins has not been compiled, and the molecular detail on UGGT:substrate interactions remains uncharacterized, it is apparent that the enzyme is highly promiscuous. Suggestions that the UGGT2 isoform (only present in higher eukayotes) is competent in re-glucosylating glycopeptides [21] may point to a duplication of the gene and evolution of two isoforms with separate pools of misfolded glycoprotein substrates. If this is the case, the “UGGT1-ome” and “UGGT2-ome” (defined as full lists of clients of UGGT1 and UGGT2, respectively [22]) would contain distinct (although possibly overlapping) sets of substrate glycoproteins. Still, how can each UGGT isoform re-glucosylate misfolded glycoproteins of a wide variety of different sizes and shapes, each substrate glycoprotein potentially presenting a unique relative orientation and distance between the site of misfold and the N-linked glycan receiving the glucose, constitutes a big open question. A second major question is the molecular basis for UGGT glycoprotein misfold recognition.

Our MD simulations of *Ct*UGGT confirm that - despite its tightly woven topology [5] - the enzyme is indeed quite flexible. More importantly, analysis of the observed MD conformational landscape for UGGT establishes for the first time the framework necessary to discuss its dynamics. The molecule’s inter-domain conformational motions can be described in terms of three simple ‘conformational coordinates’ (CCs): the relative movement between domains TRXL3 and TRXL1, resulting in the opening and closing of the cleft between them (‘clamping’, along CC1); the movement restricted to TRXL2 moving closer or further away from the relatively rigid core comprised of domains GT24-βS1-βS2-TRXL4 (‘bending’, along CC2); and the rotation of TRXL2 with respect to TRXL3 (‘twisting’, along CC3). The three motions are to some extent independent of each other, which opens the way to the cloning, expression and purification of Cys quadruple mutants such as *Ct*UGGT ^G177C/A786C,D611C/G1050C^, blocking the molecule in a clamped and bent shut conformation across the cleft and the central saddle, respectively. Such pairs of disulphide bridges forcing clamped and bent shut conformations would likely aid structural studies of mammalian UGGTs, which so far have resisted structural determination [3].

All purified UGGTs described in the literature (either recombinantly-expressed or tissue-purified) have so far revealed cleavage in the flexible linker between the folding sensor N-terminal portion and the catalytic GT24 domain (see a survey in [5]), with one study speculating large relative movements between the two portions of the UGGT molecule thanks to this flexible linker [6, 7]. In this ‘reach-and-grab’ model, UGGT would preferentially bind and re-glucosylate glycans close to the site of misfold, but would also be able to extend to re-glucosylate distal glycans in neighboring folded regions, if the GT24 were to escape the embrace of the βS1-βS2 domains [13]. Atomic force microscopy (AFM) can indeed pull the BT24 and βS1-βS2 domains apart [6, 7], but this likely constitutes mechanical denaturation, breaking the interface between these domains in a non-physiological manner. Here, we consult all the available structural evidence (namely crystal structures of full length UGGTs and their mutants, their MD trajectories and the 25 Å negative stained EM map for the complex between *Td*UGGT and an anti-*Td*UGGT Fab [6, 7]) and find no evidence suggesting separation of the βS1-βS2 and GT24 domains on either side of the cleaved flexible linker. Claims to the contrary in [6, 7] were likely due to difficulties in docking the N-terminal (PDB ID 5Y7O) and C-terminal portions (PDB ID 5H18) of *Td*UGGT separately into the negative-stain EM map, in absence of the knowledge of the intimate association between the GT24 domain and the βS1-βS2 tandem domains, observed for the first time in full-length UGGT crystal structures [5] (the *Td*UGGT study was in the last stages of the editorial process at that time).

Having concluded that UGGT’s promiscuity is not dependent on the flexible linker between the catalytic domain and the N-terminal misfold sensing region, but is likely underpinned by the motions uncovered by the MD simulations, the question remains regarding UGGT’s reported ability to survey not only folding of small- and medium-size glycoprotein monomers, but also quaternary structure of glycoprotein oligomers and larger multi-glycoprotein complexes [23] [24] [25]. The UGGT inter-domain MD movements extend beyond what was observed in the crystal structures, and indeed *in silico* modelling suggests that the extended UGGT conformations sampled by MD could accommodate glycoprotein substrates of different sizes, enabling UGGT re-glucosylation to work across a range of distances between a N-glycosylation site and a site of misfold.

Indeed, the MD conformational landscape observed in this work reaches extremely compact conformations, which may explain the activity of the enzyme against synthetic glycopeptides [16–20]. Importantly, monomeric UGGT can recognize and re-glucosylate a misfolded glycoprotein only if it can bridge the distance between a folding defect and at least one of the glycoprotein’s N-linked glycosylation sites. Thus, irrespective of the misfolded glycoprotein substrate, the finite size of the enzyme puts an upper limit to the maximum distance between a site of misfold and an N-linked glycan that monomeric UGGT can re-glucosylate on the same glycoprotein substrate (unless UGGT misfolded glycoprotein recognition is mediated by UGGT dimers/multimers – a hypothesis not supported by any data so far). The existence of this limit in turn would imply evolutionary pressure on a secreted glycoprotein sequence to develop N-glycosylations sites at accessible distances from the portions of glycoprotein that are most prone to folding difficulties (*i.e.* a folding glycoprotein ‘Achille’s heels). We propose to name “Parodi limit” [12] the maximum distance between a UGGT substrate’s site of misfold and an N-linked glycan on the same substrate that enables re-glucosylation by monomeric UGGT in response to recognition of misfold at that same site. On the basis of our *Ct*UGGT MD simulations at 300 K and on the conformational mobility of Man_9_GlcNAc_2_ N-linked glycans [26], we estimate the Parodi limit to be in the region of 60-70 Å. Functional data from UGGT-mediated re-glucosylation of a series of rigid, misfolded UGGT glycoprotein substrates, each bearing one recombinantly engineered N-linked glycosylation site at a specific distance from a single site of misfold common to all substrates in the series, would enable experimental estimation of the Parodi limit. Ideally, one such series of artificial N-linked glycosylation sites at varying distance from a single site of misfold would have to be engineered for a number of different substrate glycoprotein scaffolds, in order to minimise the dependency of the Parodi limit estimation from a given substrate series, and to estimate a standard error on that value.

When it comes to correlating UGGT interdomain conformational mobility with its activity, among the *Ct*UGGT double cysteine mutants tested so far, the *Ct*UGGT^D611C/G1050C^ mutant described in [5] is the least active in re-glucosylating urea-misfolded bovine thyroglobulin, compatible with our observation that its MD trajectory is the most severely limited one. The extra disulfide bridge engineered in this mutant joins the βS2-TRXL2 domains, therefore this mutant’s impaired activity suggests that during the enzyme:substrate encounter, a portion of misfolded thyroglobulin may be accommodated in the UGGT central saddle between the βS2 and TRXL2 domains. The *Ct*UGGT crystal structures, the cryo-EM structure [5] and our MD simulations all highlight the TRXL2 domain as the most mobile in the structure, supporting this hypothesis. The total loss of activity of the *Ct*UGGT-ΔTRXL2 construct in the re-glucosylation of urea-misfolded bovine thyroglobulin also supports a critical role for the TRXL2 domain in adjusting the size / varying the surface area of the saddle, making the bending motion a crucial one for UGGT activity against this substrate.

When it comes to the clamping motion, two *a priori* rather similar double cysteine mutants, *Ct*UGGT^S180C/T742C^ and *Ct*UGGT^G177C/A786C^, both designed to clamp the TRXL1-TRXL3 domains shut, differ significantly in their ability to re-glucosylate urea-misfolded bovine thyroglobulin, with the former mutant more active than (and the latter mutant having similar activity to) WT *Ct*UGGT. These observations point to the possibility that each misfolded glycoprotein substrate may depend to a different degree on a different subset of UGGT inter-domain conformational degrees of freedom. In the light of these data, the extent to which various portions of the UGGT structure and its motions are critical to its activity will profit from a number of re-glucosylation assays using the same set of UGGT mutants on different glycoprotein substrates.

As to the portion(s) of UGGT that mediate the binding to misfolded glycoproteins - involvement of TRXL2 and placement of substrates between this domain and the βS2-βS2-GT24 rigid domains group is the simplest hypothesis supported by the data so far. Substrate recruitment via TRXL2 movements would not require complete burial of the misfolded glycoprotein into the central saddle of the molecule: UGGT would minimally need to establish contact with the portion of substrate containing the misfold. This is plausible for relatively big substrates like transferrin (77 KDa, radius of gyration (rog) = 29.7 Å) or urea-misfolded bovine thyroglobulin (670 KDa, a long snake-like chain of eleven 60 AA compact domains, no structure available). For smaller substrates, like glycopeptides or synthetic fluorescent probes, closer UGGT conformations, bringing TRXL2 towards βS2 and the GT24 domain across the central saddle, may be needed. Apart from TRLX2, other untested UGGT regions potentially harboring exposed hydrophobic patches are the *Ct*UGGT TRXL4 disordered region 243–285; the flexible linker around the endo-proteolysis site between the βS2 and GT24 domains (*Ct*UGGT 1153-1195), and the residues between the last helix and the ER retrieval motif at the C-terminus (*Ct*UGGT 1474-1510) [5]. Again, structural and functional data from a range of UGGT mutants and glycoprotein substrates will be required to further test these hypotheses and fully dissect the UGGT structure-function relationship.

Little if any structural changes are visible in the GT24-βS1-βS2 portion of the molecule across the ensemble of available structures and during MD simulations, the beta sandwiches largely shielding the catalytic domain from the TRXL2 domains movements. The footprint of the N-linked glycan is likely a conserved grooved on the surface of the GT24 domain - residues *Ct*UGGT 1276-1280, 1338-1346, 1392-1402 [5] - but no mutagenesis has probed this hypothesis yet. Glucosyltransferase domains of the GT24 family have been shown to possess conformational plasticity, requiring structural changes during substrate recognition and catalysis [27, 28]. Based on the MD simulations, the interface between domains TRXL3 and TRXL4 could act as a hinge communicate changes across the central part of the molecule, but in absence of structural information of UGGT:substrate complexes, it is not possible to rule out local changes in the catalytic domain induced by the N-linked glycan docking onto its surface, nor allosteric changes communicated to the catalytic domain by the binding of the glycoprotein substrate to the N-terminal misfold-sensing region. Yet, the simplest model of UGGT activity compatible with the data observed so far remains one in which a substrate glycoprotein binding to the N-terminal misfold-sensing region allows prolonged association between enzyme and substrate, increasing the avidity of the catalytic domain for the N-linked glycan. In this model, the GT24 domain of UGGT undergoes no major structural changes when accommodating the Man_9_GlcNAc_2_ N-linked glycan, and no allostery is at play between N- and C-terminal portions of the enzyme.

The molecular forces supporting UGGT-mediated glycoprotein misfold recognition have been generally hypothesized to be hydrophobic/hydrophobic interactions [29]. Our observation that the UGGT TRXL2 domain surface facing the central saddle bears distinct patches of hydrophobic residues that are conserved across UGGT1 and UGGT2 sequences [5] supports such models of misfold recognition. The fact that UGGT bears de-mannosylated glycans - a hallmark of ER associated degradation (ERAD) [30, 31] - is also compatible with the hypothesis that UGGT may recognise misfolded glycoproteins *via* an intrinsically misfolded domain (“it takes one to know one” [22]), as observed for the mouse ERAD checkpoint mannosidase, which also preferentially acts on misfolded glycoproteins and has been proven to undergo constant ERAD degradation [32]. The hypothesis that UGGT works by having evolved an intrinsically misfolded portion, with which the enzyme would interact with substrate glycoprotein misfolded regions, would in turn imply that UGGT may re-glucosylate itself, eventually being subjected to ERAD de-mannosylation and degradation. The associated biochemical costs of UGGT self-reglucosyation and ERAD de-mannosylation/degradation may be the price eukaryotic cells evolved to pay in order to afford UGGT as a universal glycoprotein misfolding checkpoint. *In vitro* and *in cellula* experiments to test these ideas are in progress.

## Experimental Procedures

Cloning, expression and purification of full-length *Ct*UGGT are described in [5].

### Cloning

#### Cloning of *Ct*UGGT^G177C/A786C^ and *Ct*UGGT^V178C/A786C^

Mutation of the *Ct*UGGT into *Ct*UGGT^A786C^ was effected starting from the gene of *Ct*UGGT inserted in Litmus28i as follows: 12.5 µl of Q5® Hot Start High-Fidelity 2X Master Mix (New England Biolabs) were added to 1.25 µl of each forward and reverse primers (A786C_F and A786C_R in SI Appendix S3) at 10 µM, 1 µl of *Ct*UGGT:Litmus28i DNA at 1 ng/µL and 9 µl of nuclease-free water, obtaining a 25 µl final volume. PCR amplification: step 1: 98 °C for 30 s; step 2: 98 °C for 10 s; step 3: 60 °C for 20 s; step 4: 72 °C for 135 s. Steps 2-4 were repeated 25 times; step 5: 72 °C for 2 minutes. Kinase, Ligase & DpnI (KLD) treatment deleted the parental fragments present in the mixture: 1 µL of PCR product was mixed with 5 µL of 2X KLD Reaction Buffer, 1 µL of 10X KLD Enzyme Mix (both from New England Biolabs) and 3 µL of nuclease-free water. The mixture was incubated at room temperature for 5 minutes. The KLD reaction mixture was used to transform *E. coli* DH-5α chemically competent cells using the following protocol: 5 µl of KLD reaction mix were added to a tube of thawed New England BioLabs DH-5α competent *E. coli* cells on ice, and mixed gently for a few seconds; after the transformation, the bacteria were incubated on ice for 30 minutes, heat shocked at 42 °C for 30 seconds and incubated on ice again for 5 minutes. 950 µl of SOC media were added to a final volume of 1 ml and the mixture was incubated for 1 hour at 37 °C with gentle shaking at 200/300 rpm. 100 µl of the bacteria were spread onto a pre-warmed (37 °C) LB agar culture plate containing carbenicillin (0.1 mg/mL). The plate was incubated at 37 °C overnight. Colony-PCR was performed on DNA from various colonies (using T7_F and T7_R primers, see Appendix) and the DNA obtained was loaded on a 1% w/v agarose gel and run for 50 minutes at 150 V. Analysis of this gel allowed identification of colonies with amplified DNA of the appropriate size, which were used to inoculate 5 ml LB supplemented with 0.1 mg/mL carbenicillin. This was used to make glycerol stocks, mixing 16% glycerol with 84% bacteria and freezing and storing at −80 °C. The DNA was mini-prepped (Qiagen) and after sequencing (with primer *Ct*UGGT_401_800_F, SI Appendix Table S3), a glycerol stock of one of the previously described colonies was used to inoculate 5 ml LB supplemented with 0.1 mg/mL carbenicillin and incubated over night at 37 °C. This culture was then used to inoculate 200 ml LB supplemented with 0.1 mg/mL carbenicillin and left to incubate at 37 °C, shaking at 110 rpm. Upon reaching an OD_600nm_ of 2.0, the cells were spun down at 3320xg for 18 minutes. The pellets were resuspended, and the DNA purified using a Maxiprep kit (Qiagen), following the recommended protocol to obtain 3 mL of *Ct*UGGT^A786C^:Litmus28i plasmid DNA at 400 ng/ml and 3 mL of *Ct*UGGT^A786C^:Litmus28i plasmid DNA at 400 ng/µL.

To obtain the double mutant *Ct*UGGT^G177C/A786C^ (*Ct*UGGT^V178C/A786C^), the second mutation G177C (V178C) was introduced starting from the gene of *Ct*UGGT^A786C^ in Litmus28i as follows: 12.5 µl of Q5® Hot Start High-Fidelity 2X Master Mix (New England Biolabs) were added to 1.25 µl of each forward and reverse primers (G177C: primers G177C _F and G177C _R; V178C: primers V178C _F and V178C_R, see SI Appendix Table S3) at 10 µM, 1 µl of *Ct*UGGT^A786C^:Litmus28i DNA at 1 ng/µL and 9 µl of nuclease-free water, obtaining a 25 µl final volume. PCR protocol: step 1: 98 °C for 30 seconds; step 2: 98 °C for 10 seconds; step 3: *Ct*UGGT^G177C/A786C^: 66 °C for 30 seconds (*Ct*UGGT^V178C/A786C^: 60 °C for 20 seconds); step 4: 72 °C for 135 seconds; steps 2-4 were repeated 25 times. Step 5: 72 °C for 2 minutes. After KLD treatment (see above) *E. coli* DH-5α chemically competent cells were transformed with the DNA as described previously. Colony-PCR was performed on DNA from various colonies (using T7_F and T7_R primers, see SI Appendix Table S3) and the DNA obtained was loaded on a 1% w/v agarose gel and run for 50 minutes at 150 V. Analysis of this gel allowed identification of colonies with amplified DNA of the appropriate size, which were used to inoculate 5 ml LB supplemented with 0.1 mg/mL carbenicillin. This was used to make glycerol stocks (see above). The DNA was mini-prepped (Qiagen), sequenced with primers X_F and X_R and maxiprepped (see above) to obtain 3 mL of *Ct*UGGT^G177C/A786C^:Litmus28i plasmid DNA at 700 ng/µL and 3 mL of *Ct*UGGT^V178C/A786C^:Litmus28i plasmid DNA at 500 ng/µL.

The *Ct*UGGT^G177C/A786C^ and *Ct*UGGT^V178C/A786C^ inserts in Litmus 28i were separately cloned into the pHLsec vector [33] to contain a hexa-His Tag at the C-terminus. DNA for pHLsec was linearised using AgeI and KpnI restriction enzymes at 37 °C for 16 hours in Cutsmart™ buffer (New England Biolabs). The restriction digest was then run on a 0.8% w/v agarose gel at 150 V for 1 hour. The linearised vector was cut from the gel and purified with a QIAquick gel extraction kit (Qiagen). PCR was performed on the *Ct*UGGT^G177C/A786C^ insert in Litmus 28i (or *Ct*UGGT^V178C/A786C^ insert in Litmus 28i): 1 µl (1 ng/µl) added at 25 µl of Q5® Hot Start High-Fidelity 2X Master Mix (New England Biolabs), 2.5 µl of each forward and reverse primers (pHLsec_*Ct*UGGT_F and pHLsec_*Ct*UGGT_R, see SI Appendix Table S3) and 19 µl of nuclease-free water. PCR protocol: step 1: 98 °C for 30 seconds; step 2: 98 °C for 10 seconds; step 3: 62 °C for 30 seconds; step 4: 70 °C for 150 seconds; step 2-4 were repeated 35 times; step 5: 72 °C for 2 minutes. The PCR products were run on a 0.8% w/v agarose gel at 150 V for 1 hour and the amplified insert was cut from the gel and purified with the same QIAquick gel extraction kit. A Gibson Assembly was then performed using the gel-purified PCR-amplified *Ct*UGGT^G177C/A786C^ (or *Ct*UGGT^V178C/A786C^) insert mixed with gel-purified linearised pHLsec at a ratio of 3:1 with NEBuilder® HiFi DNA Assembly Master Mix (New England Biolabs) using suggested protocol, for 1 hour at 50 °C. 2 µl of this ligation product was added to 50 µl XL10-Gold Ultracompetent cells (Agilent), following the transformation guideline protocol. The cells were then plated on 0.1 mg/mL carbenicillin agar plates and incubated over night at 37 °C.

Colony-PCR was performed on DNA from various colonies (using pHLsec_F and pHLsec_R primers, see SI Appendix Table S3) and the DNA obtained was run on a 1% w/v agarose gel for 50 minutes at 150 V. Analysis of this gel allowed identification of colonies with amplified DNA of the appropriate size, which were miniprepped, glycerol-stocked (sequenced with primers *Ct*UGGT_400_1_R and *Ct*UGGT_401_800_F see SI Appendix Table S3), and maxiprepped as described earlier, to obtain 3 mL of *Ct*UGGT^G177C/A786C^:pHLsec plasmid DNA at 300 ng/µL and 3 mL of *Ct*UGGT^V178C/A786C^:pHLsec plasmid DNA at 700 ng/µL.

#### Cloning of *Ct*UGGT^S180C/T742C^

Mutation of the *Ct*UGGT into *Ct*UGGT^T742C^ was effected starting from the gene of *Ct*UGGT inserted in Litmus28i as follows: 12.5 µl of Q5® Hot Start High-Fidelity 2X Master Mix (New England Biolabs) were added to 1.25 µl of each forward and reverse primers (T742C_F and T742R in SI Appendix S3) at 10 µM, 1 µl of *Ct*UGGT:Litmus28i DNA at 1 ng/µL and 9 µl of nuclease-free water, obtaining a 25 µl final volume. PCR amplification: step 1: 98 °C for 30 s; step 2: 98 °C for 10 s; step 3: 59 °C for 20 s; step 4: 72 °C for 135 s. Steps 2-4 were repeated 25 times; step 5: 72 °C for 2 minutes. Kinase, Ligase & DpnI (KLD) treatment deleted the parental fragments present in the mixture: 1 µL of PCR product was mixed with 5 µL of 2X KLD Reaction Buffer, 1 µL of 10X KLD Enzyme Mix (both from New England Biolabs) and 3 µL of nuclease-free water. The mixture was incubated at room temperature for 5 minutes. The KLD reaction mixture was used to transform *E. coli* DH-5α chemically competent cells using the following protocol: 5 µl of KLD reaction mix were added to a tube of thawed New England BioLabs DH-5α competent *E. coli* cells on ice, and mixed gently for a few seconds; after the transformation, the bacteria were incubated on ice for 30 minutes, heat shocked at 42 °C for 30 seconds and incubated on ice again for 5 minutes. 950 µl of SOC media were added to a final volume of 1 ml and the mixture was incubated for 1 hour at 37 °C with gentle shaking at 200/300 rpm. 100 µl of the bacteria were spread onto a pre-warmed (37 °C) LB agar culture plate containing carbenicillin (0.1 mg/mL). The plate was incubated at 37 °C overnight. Colony-PCR was performed on DNA from various colonies (using T7_F and T7_R primers, see Appendix) and the DNA obtained was loaded on a 1% w/v agarose gel and run for 50 minutes at 150 V. Analysis of this gel allowed identification of colonies with amplified DNA of the appropriate size, which were used to inoculate 5 ml LB supplemented with 0.1 mg/mL carbenicillin. This was used to make glycerol stocks (mixing 16% glycerol with 84% bacteria and freezing and storing at −80 °C. The DNA was mini-prepped (Qiagen) and after sequencing (with primer *Ct*UGGT_401_800_F, SI Appendix Table S3), a glycerol stock of one of the previously described colonies was used to inoculate 5 ml LB supplemented with 0.1 mg/mL carbenicillin and incubated over night at 37 °C. This culture was then used to inoculate 200 ml LB supplemented with 0.1 mg/mL carbenicillin and left to incubate at 37 °C, shaking at 110 rpm. Upon reaching an OD_600nm_ of 2.0, the cells were spun down at 3320xg for 18 minutes. The pellets were resuspended, and the DNA purified using a Maxiprep kit (Qiagen), following the recommended protocol to obtain 3 mL of *Ct*UGGT^A742C^:Litmus28i plasmid DNA at 500 ng/µL.

To obtain the double mutant *Ct*UGGT^S180C/A742C^, the second mutation S180C was introduced starting from the gene of *Ct*UGGT^A742C^ in Litmus28i as follows: 12.5 µl of Q5® Hot Start High-Fidelity 2X Master Mix (New England Biolabs) were added to 1.25 µl of each forward and reverse primers (primers S180C_F and S180C_R, see SI Appendix Table S3) at 10 µM, 1 µl of *Ct*UGGT^A742C^:Litmus28i DNA at 1 ng/µL and 9 µl of nuclease-free water, obtaining a 25 µl final volume. PCR protocol: step 1: 98 °C for 30 seconds; step 2: 98 °C for 10 seconds; step 3: 68 °C for 30 seconds; step 4: 72 °C for 135 seconds; steps 2-4 were repeated 25 times. Step 5: 72 °C for 2 minutes. After KLD treatment (see above) *E. coli* DH-5α chemically competent cells were transformed with the DNA as described previously. Colony-PCR was performed on DNA from various colonies (using T7_F and T7_R primers, see SI Appendix Table S3) and the DNA obtained was loaded on a 1% w/v agarose gel and run for 50 minutes at 150 V. Analysis of this gel allowed identification of colonies with amplified DNA of the appropriate size, which were used to inoculate 5 ml LB supplemented with 0.1 mg/mL carbenicillin. This was used to make glycerol stocks (see above). The DNA was mini-prepped (Qiagen), sequenced with primers X_F and X_R and maxiprepped (see above) to obtain 3 mL of *Ct*UGGT^S180C/T742C^:Litmus28i plasmid DNA at 500 ng/µl.

The *Ct*UGGT^S180C/T742C^ insert in Litmus 28i was cloned into the pHLsec vector [33] to contain a hexa-His Tag at the C-terminus. DNA for pHLsec was linearised and gel-purified as described above for the *Ct*UGGT^G177C/T786C^ and *Ct*UGGT^V178C/T786C^ double mutants. PCR was performed on the *Ct*UGGT^S180C/A742C^ insert in Litmus 28i: 1 µl (1 ng/µl) added at 25 µl of Q5® Hot Start High-Fidelity 2X Master Mix (New England Biolabs), 2.5 µl of each forward and reverse primers (pHLsec_*Ct*UGGT_F and pHLsec_*Ct*UGGT_R, see SI Appendix Table S3) and 19 µl of nuclease-free water. PCR protocol: step 1: 98 °C for 30 seconds; step 2: 98 °C for 10 seconds; step 3: 62 °C for 30 seconds; step 4: 70 °C for 150 seconds; step 2-4 were repeated 35 times; step 5: 72 °C for 2 minutes. The PCR products were run on a 0.8% w/v agarose gel at 150 V for 1 hour and the amplified insert was cut from the gel and purified with the same QIAquick gel extraction kit. A Gibson Assembly was then performed using the gel-purified PCR-amplified *Ct*UGGT^S180C/A742C^ insert mixed with gel-purified linearised pHLsec at a ratio of 3:1 with NEBuilder® HiFi DNA Assembly Master Mix (New England Biolabs) using suggested protocol, for 1 hour at 50 °C. 2 µl of this ligation product was added to 50 µl XL10-Gold Ultracompetent cells (Agilent), following the transformation guideline protocol. The cells were then plated on 0.1 mg/mL carbenicillin agar plates and incubated over night at 37 °C.

Colony-PCR was performed on DNA from various colonies (using pHLsec_F and pHLsec_R primers, see SI Appendix Table S3) and the DNA obtained was run on a 1% w/v agarose gel for 50 minutes at 150 V. Analysis of this gel allowed identification of colonies with amplified DNA of the appropriate size, which were miniprepped, glycerol-stocked (sequenced with primers *Ct*UGGT_400_1_R and *Ct*UGGT_401_800_F see SI Appendix Table S3), and maxiprepped as described earlier, to obtain 3 mL of *Ct*UGGT^S180C/A742C^:pHLsec plasmid DNA at 700 ng/µL.

#### Cloning of *Ct*UGGT-ΔTRXL1

The *Ct*UGGT-ΔTRXL1 construct lacks residues *Ct*UGGT 42-224. The deletion of the *Ct*UGGT TRXL1 domain was performed starting from the gene of *Ct*UGGT in Litmus28i as follows: 12.5 µl of Q5® Hot Start High-Fidelity 2X Master Mix (New England Biolabs) were added to 1.25 µl of each forward and reverse primers at 10 µM (primers Δ1_F and Δ1_R, see SI Appendix Table S3), 1 µl of *Ct*UGGT DNA 1 ng/µL and 9 µl of nuclease-free water, obtaining a 25 µl final volume; PCR protocol: step 1: 98 °C for 30 seconds; step 2: 98 °C for 10 seconds; step 3: 65 °C for 20 seconds; step 4: 72 °C for 130 seconds; step 2-4 were repeated 25 times; step 5: 72 °C for 2 minutes. KPN treatment and *E. coli* XL10-Gold Ultracompetent cells (Agilent) transformation and plating as described before. Colony-PCR was performed on DNA from various colonies (using T7_F and T7_R primers, see SI Appendix Table S3) and the DNA obtained was run on a 1% w/v agarose gel for 50 minutes at 150 V. Analysis of this gel allowed identification of colonies with amplified DNA of the appropriate size, which were used to miniprep the DNA, sequenced with primer pHLsec_F, see SI Appendix Table S3) and maxiprepped to obtain 3 mL of *Ct*UGGT-ΔTRXL1:Litmus28i plasmid DNA at 400 ng/µl. The *Ct*UGGT-ΔTRXL1 insert in Litmus 28i was cloned into the pHL-sec vector to contain a hexa-His Tag at the C-terminus as described before for the double Cys mutants to obtain 3 mL *Ct*UGGT-ΔTRXL1:pHL-sec plasmid DNA at 500 ng/µl.

#### Cloning of *Ct*UGGT-ΔTRXL2

the *Ct*UGGT-ΔTRXL2 construct lacks residues *Ct*UGGT 417-650. The deletion of the *Ct*UGGT TRXL2 domain was performed starting from the gene of *Ct*UGGT in Litmus28 (an optimal vector for mutagenesis experiments) as follows: 25 µL of 2X Q5 high fidelity master mix (New England Biolabs) were added to 1.25 µL of 10 µM forward and reverse primers Δ2_F and Δ2_R (see SI Appendix Table S3), 1 µL of 20 ng/µL *Ct*UGGT DNA and 9 µL of water; PCR protocol: step 1: 98 °C for 30 seconds; step 2: 98 °C for 10 seconds; step 3: 64 °C for 20 seconds; step 4: 72 °C for 180 seconds; step 2-5 were repeated 25 times; step 6: 72 °C for 2 minutes. Kinase, Ligase & DpnI (KLD) treatment was made to delete the parental fragments present in the mixture. 1 µL of PCR product was mixed with 5 µL of 2X KLD Reaction Buffer, 1 µL of 10X KLD Enzyme Mix and 3 µL of water. The mixture was incubated at room temperature for 5 minutes.

The CtUGGT-ΔTRXL2 insert in Litmus 28i was cloned into the pHLsec vector [33] in frame to contain a hexa-His Tag at the C-terminus. DNA for pHLsec was linearised using AgeI and KpnI restriction enzymes at 37°C for 16 hours in Cutsmart™ buffer (NEB). The restriction digest was then run on a 0.8% w/v agarose gel at 150V for 1 hour. The linearised vector was cut from the gel and purified with a QIAquick gel extraction kit (Qiagen). PCR was performed on the CtUGGT-ΔTRXL2 insert in Litmus 28i using Q5 MasterMix (NEB) and a primer annealing temperature of 70°C. The primers used were *Ct*UGGT_pHLsec_F and *Ct*UGGT_phLsec_R (see SI Appendix Table S3). The PCR products were run on a 0.8% w/v agarose gel at 150V for 1 hour and the amplified insert was cut from the gel and purified with the same QIAquick gel extraction kit. A Gibson Assembly was then performed using the gel-purified PCR-amplified CtUGGT-ΔTRXL2 insert mixed with gel-purified linearised pHLsec at a ratio of 3:1 with HiFi Q5 MasterMix (NEB) for 1 hour at 50°C. 5µl of this ligation product was added to 50µl NEB5α competent *E. coli* (High Efficiency) cells (NEB), followed by the guideline transformation protocol. The cells were then plated on 0.1 mg/mL carbenicillin agar plates and incubated for 16 hours at 37°C. Colony-PCR was performed on DNA from various colonies (using the same primers as above) and the DNA obtained was run on a 0.8% w/v agarose gel for 1 hour at 150V. Analysis of this gel allowed identification of colonies with amplified DNA of the appropriate size, which were used to inoculate 5ml LB supplemented with 0.1 mg/mL carbenicillin. This was used to make glycerol stocks. The DNA was mini-prepped (Qiagen) and after sequencing, a glycerol stock of one of the previously described colonies was used to inoculate 5ml LB supplemented with 0.1 mg/mL carbenicillin and incubated for 16 hours at 37°C. This culture was then used to inoculate 200ml LB supplemented with 0.1 mg/mL carbenicillin and left to incubate at 37°C, shaking at 110rpm. Upon reaching an OD_600nm_ of 2, the cells were spun down at 3320xg for 18 minutes. The pellets were resuspended and the DNA purified using a Maxiprep kit (Qiagen), following the recommended protocol to obtain 3ml *Ct*UGGTΔTRXL2:pHLsec plasmid DNA at 300 ng/µL.

#### Cloning of *Ct*UGGT-ΔTRXL3

The *Ct*UGGT-ΔTRXL3 construct lacks residues *Ct*UGGT 666-870. The deletion of the *Ct*UGGT TRXL3 domain was performed starting from the gene of *Ct*UGGT in Litmus28i as follows: 12.5 µl of Q5® Hot Start High-Fidelity 2X Master Mix (New England Biolabs) were added to 1.25 µl of each forward and reverse primers at 10 µM (primers Δ3_F and Δ3_R, see SI Appendix Table S3), 1 µl of *Ct*UGGT DNA 1 ng/µL and 9 µl of nuclease-free water, obtaining a 25 µl final volume; PCR protocol: step 1: 98 °C for 30 seconds; step 2: 98 °C for 10 seconds; step 3: 62 °C for 20 seconds; step 4: 72 °C for 130 seconds; step 2-4 were repeated 25 times; step 5: 72 °C for 2 minutes. KPN treatment and *E. coli* XL10-Gold Ultracompetent cells (Agilent) transformation and plating as described before. Colony-PCR was performed on DNA from various colonies (using T7_F and T7_R primers, see SI Appendix Table S3) and the DNA obtained was run on a 1% w/v agarose gel for 50 minutes at 150 V. Analysis of this gel allowed identification of colonies with amplified DNA of the appropriate size, which were used to miniprep the DNA, sequenced with primer *Ct*UGGT_401_800_F, see SI Appendix Table S3) and maxiprepped to obtain 3 mL of *Ct*UGGT-ΔTRXL1:Litmus28i plasmid DNA at 500 ng/µl. The *Ct*UGGT-ΔTRXL3 insert in Litmus 28i was cloned into the pHL-sec vector to contain a hexa-His Tag at the C-terminus as described before for the double Cys mutants to obtain 3 mL *Ct*UGGT-ΔTRXL3:pHL-sec plasmid DNA at 800 ng/µl.

### Protein Production

All transfections were carried out as follows, except where otherwose indicated. Human epithelial kidney FreeStyle 293F cells (ThermoFisher Scientific) at 10^6^ cells/mL suspended in FreeStyle 293 Media (ThermoFisher Scientific) were transfected using the FreeStyle 293 expression system (ThermoFisher Scientific). For a 50mL culture: 62.5 µL of FreeStyle MAX transfection reagent (ThermoFisher Scientific) and 62.5 µg of plasmid DNA were each separately diluted to 1 mL with OptiPRO SFM reagent (ThermoFisher Scientific) then mixed, incubated for 10 min at room temperature and finally added to the cell suspension. Transfected cells were left shaking in 500 mL Erlenmeyer flasks with 0.2 µm vent caps (Corning) shaking at 135 revolutions per min (rpm) in a 37 °C incubator kept at 8% CO2. The HEK-293 cells at 10^6^ cells/litre were transfected by mixing the *Ct*UGGTΔTRXL2 pHLsec plasmid DNA, and Lipofectamine Transfection Reagent (Thermofisher), with Optipro (Thermofisher) in separate tubes, before mixing the two tubes, leaving them for 10 minutes at room temperature and then adding to the cells. The cells were left shaking in 500 mL Erlenmeyer flasks with 0.2µm vent caps at 135rpm for three days at 37°C, 8% CO_2_.

#### *Ct*UGGT_Kif_

to express *Ct*UGGT_Kif_, 300 mL human epithelial kidney FreeStyle 293 (HEK293F) cells at 10^6^ cells/mL, suspended in GIBCO FreeStyle 293 Media, supplemented with 5 µM kifunensine (Cayman Chemical Company), were transfected using the FreeStyle MAX 293 expression system, according to manifacturer instructions. >90 % cell viability was confirmed by trypan blue exclusion. A volume of 375 µL of FreeStyle MAX transfection reagent and 300 µg plasmid DNA *Ct*UGGT:pHLsec in 375 µL of water were each separately diluted to 6 mL with OptiPRO SFM reagent, then mixed and incubated for 7 minutes at room temperature. The mixture was split evenly between two cultures, each containing total 150 mL of HEK293F cells at a density of 10^6^ cells/mL. Transfected cells were left shaking at 135 revolutions per minute in 0.5 L Erlenmeyer flasks with 0.2 µm vent caps (Corning), in an incubator at 37 °C with 5 % CO_2_ present, for 6 days.

The HEK293F cells were separated from the supernatant by centrifugation for 15 minutes at 4 °C and 3,000 g. The supernatant was made to contain to 1x phosphate buffered saline (PBS) and 5 mM imidazole by adding appropriate stock solution volumes, and pH adjusted to 7.4 by adding a few drops of 2 M NaOH, before vacuum filtration through a 0.45 µm filter and application onto a 1 mL HisTrap HP Ni IMAC column (GE Healthcare) equilibrated against binding buffer: 1x PBS, 5 mM imidazole, pH adjusted to 7.4 with a few drops of 2 M NaOH. The column was washed with 20 column volumes (cV) buffer A and protein eluted with a linear gradient over 20 cV from 0 % to 100 % of elution buffer B: 1x PBS, 400 mM imidazole, pH adjusted to 7.4 with 2 M NaOH. Peak fractions were pooled and concentrated using PES membrane, 50 kDa MW cutoff centrifugal ultrafiltration devices (Sartorius), to a volume of 5 mL. Concentrated *Ct*UGGT_Kif_ sample was applied to a HiLoad Superdex 200 16/60 size exclusion chromatography (SEC) column (GE Healthcare) equilibrated against SEC buffer: 20 mM NaHEPES, 150 mM NaCl. Peak fractions were pooled and concentrated as before, protein concentration measured by loading 1.5 µL of sample on a NanoDrop 1000 spectrophotometer (Thermo Scientific). The calculated ε_280_ was then used to estimate the protein concentration. The final concentration of protein (1 mL volume) was 7.24 mg/mL (A_280_ = 8.18). *Ct*UGGT_Kif_ protein aliquots were frozen in liquid N_2_ and stored at −80 °C. SDS-PAGE of SEC fractions was used to assess purity. All chromatography was at 1 mL/min flow rate on ÄKTA Pure (room temperature) or ÄKTA Start (4 °C) systems (GE Healthcare).

#### *Ct*UGGT-ΔTRXL2

Three days after transfection, the 150 mL of HEK293F cells’ supernatant was spun for half an hour at 1,000g, made 1xPBS by addition of 10xPBS stock, brought to pH 7.6 by addition of a few drops of NaOH 10M, filtered with a 0.22 µm filter and and loaded onto a 5ml His Trap (GE Life Sciences) equilibrated and washed with ‘Buffer A’ (PBS pH 7.5), and then eluted with an increasing gradient of ‘Buffer B’ (PBS pH 7.5, 500mM imidazole) over 20 column volumes at 2 mL/min. Elution samples were analysed using SDS-PAGE and the protein-containing eluate fractions were pooled and spin-concentrated to 10ml on a 100,000 MXCO spin concentrator, before being injected in two 5ml batches onto a S200 16/60 gel filtration column (GE Life Sciences) which had been equilibrated with 20mM HEPES pH 7.2, 120mM NaCl. Elution fractions containing the desired protein were then pooled based on SDS-PAGE analysis.

#### *Ct*UGGT^S180C/T742C^

transfection and expression in HEK293F cells was carried out as described at the beginning of this section. Four days after transfection, the 400 mL of HEK293F cells’ supernatant was spun for half an hour at 1,000g, made 1xPBS by addition of 10xPBS stock, brought to pH 7.6 by addition of a few drops of NaOH 10M, filtered with a 0.22 µm filter and loaded onto a 5ml His Trap (GE Life Sciences) equilibrated and washed with ‘Buffer A’ (PBS pH 7.5), and then eluted with an increasing gradient of ‘Buffer B’ (PBS pH 7.5, 500mM imidazole) over 20 column volumes at 2 mL/min. Elution samples were analysed using SDS-PAGE and the protein was in the flow-through of the column - having apparently failed to bind to the nickel column. The 660 mL of flow-through was re-filtered through a 1 µm filter, then a 0.45 µm filter, then a 0.22 µm filter. It was diluted to 1 L with H_2_O and pH’ed to pH 8.5 with NaOH, and bound to a HiPrep Q HP 16/60 anion exchange column equilibrated in Buffer A: K_2_HPO_4_/KH_2_PO_4_ 20 mM pH 8.5, flowing at 4 mL/min. The column turned pink probably because the pH indicator from the HEK293F cells medium is anionic at pH 8.5. The column was washed with about 250 mL of A and the protein eluted using buffer B, made by making buffer A 1 M NaCl, with the following three steps: 1. 3.5 CV of 25% buffer B; 2. 3.5 CV of 50% buffer B; 3. 3.5 CV of 100% buffer B; 15 mL fractions were collected. Protein containing fractions were pooled and the 30 mL sample concentrated to 5 mL, then exchanged to 20 mM MES pH 6.5, 50 mM NaCl in two 150 KDa MWCO spin concentrators. The 5 mL of *Ct*UGGT^S180C/T742C^ in 20 mM MES pH 6.5, 50 mM NaCl were injected onto a S200 16/60 size exclusion chromatography column equilibrated in the same buffer, and run at 1 mL min, collecting 1.5 mL fractions. Protein containing fractions were pooled and concentrated to V=800 µL OD_280_=19.00 (6.28 mg/ml) and the sample stored at 4 °C.

#### *Ct*UGGT^G177C/A786C^

A 200 ml volume of HEK293F cells culture was transfected with *Ct*UGGT^S177C/A786C^:pHLsec plasmid DNA and left expressing for 4 days, the supernatant processed as described previously for *Ct*UGGT^S180C/T742C^, and run on a 5 ml HisTrap HP column (GE Life Sciences). Protein containing fractions (as detected by SDS-PAGE) were pooled and concentrated in a 50 KDa MWCO centrifugal concentrator before loading on a Superdex 200 16/60 column for size exclusion chromatography. Eluted fractions were analysed by SDS-PAGE. Protein containing fractions were pooled and concentrated as before, flash-frozen in liquid nitrogen in 100 µl aliquots and stored at −80 °C. 0.8 ml at 7.91 mg/ml of *Ct*UGGT^G177C/A786C^ were obtained.

#### *Ct*UGGT-ΔTRXL1 and *Ct*UGGT-ΔTRXL3

Transfection, expression and purification followed the protocol described for *Ct*UGGT^G177C/A786C^. After size-exclusion chromatography, 0.1 ml at 0.4 mg/ml of *Ct*UGGT-ΔTRXL1 and 0.2 ml at 12.80 mg/ml of *Ct*UGGT-ΔTRXL3 were obtained.

### Protein Crystallization

#### *Ct*UGGT_Kif_

A *Ct*UGGT_Kif_ crystal grew at 18 °C in condition 34 of the MORPHEUS screen [34] [0.09 M NPS, *aka* 0.03M Sodium nitrate, 0.03 Sodium phosphate dibasic, 0.03M Ammonium sulfate; 0.1M Buffer System 3, aka Tris Bicine pH 8.5; 8.530% v/v Precipitant Mix 2, aka 40% v/v Glycerol, 20% w/v PEG 4000] mixed in protein:mother liquor ratio 100 nL:100 nL. The crystal grew at 18 °C and it was flash-cooled in liquid N_2_. X-ray diffraction was collected at I03@Diamond.

#### *Ct*UGGT-ΔTRXL2

a *Ct*UGGT-ΔTRXL2 crystal grew at 18 °C from protein concentrated to 6.5 mg/mL and mixed in 133:67 nL protein:mother liquor ratio with solution 2 of the JCSG+ crystallisation screen in a sitting drop: 0.1M Sodium citrate pH 5.5, 20% w/v PEG 3,000. The crystal was cryo-protected with 20% glycerol in mother liquor and cryo-cooled with liquid nitrogen.

#### *Ct*UGGT^S180C/T742C^

A *Ct*UGGT^S180C/T742C^ crystal grew from protein at OD_280_=7.29 in HEPES 20 mM pH 6.5, 50 mM NaCl, 5 mM UDP-Glc, 1 mM CaCl_2_ mixed in protein:mother liquor ratio 100 nL:100 nL with condition 57 of the MORPHEUS2 screen [35] (2 mM Lanthanides, 0.1 M Buffer System 6 (1.0M, pH 8.5 at 20 °C, Gly-Gly, AMPD), 36 % v/v Precipitant Mix 5 (30% w/v PEG 3000, 40% v/v 1, 2, 4-Butanetriol, 2% w/v NDSB 256)). The crystal grew between day 57 and day 71, at 18 °C. The crystal was flash-cooled in liquid N_2_. X-ray diffraction data from the trigonal P3_2_12 and the orthorhombic P2_1_2_1_2_1_ *Ct*UGGT^S180C/T742C^ crystals were collected on beamlines I24 and I03 @Diamond, respectively.

#### *Ct*UGGT^G177C/A786C^

Initial *Ct*UGGT^G177C/A786C^ crystals grew from a solution of mother liquor: 16.54% w/v PEG 4,000, 0.03 M citric acid pH 5.3, 0.07 M citric Acid pH 6.0, 12.75% v/v isopropanol. The crystals gave very low resolution diffraction and it was decided to dehydrate them by re-equilibrating the crystallization drop against a PEG 6,000-containing mother liquor reservoir: 13 µL of mother liquor were taken out of the 50 µL in the reservoir, replaced with 13 µL of a solution of 50% w/V PEG 6,000 in mother liquor, and the plate re-sealed. After undergoing dehydration for a week, one crystal was flash frozen in liquid N_2_ for data collection.

### UGGT-mediated of urea-misfolded bovine thryoglobulin

Bovine thyroglobulin (Sigma-Aldrich) was denatured with urea following the protocol by Trombetta *et al* [11]. Each reaction mixture contained 100 µg of urea-misfolded bovine thyroglobulin, 86 µM UDP-Glucose, 8.6 mM CaCl_2_, 8.6 mM Tris-HCl pH 8.0 and 45 pmol of *Ct*UGGT enzyme (either wild-type, or one of the double Cys mutants). The reaction mixtures were set up at 37 °C. Each reaction was 70 µL to start with, in triplicate. 10 µL aliquots were taken at each time point (5’, 15’, 30’, 1 h, 2 h and O/N), and the glucosylation quenched by addition to each 10 µL aliquot of 1 µL of PNAGaseF denaturing buffer, then heating for 10 min at 90 °C. Then 5 µL of 10X PNGase glycobuffer 2 (NEB), 5 µL of NP40 10%, 1 µL of PNGase F at 1 mg/mL and 27 µL of water were added to each sample for the overnight digestion with PNGase F. The *N*-linked glycan were labelled with anthranilic acid (2-AA) (Sigma-Aldrich), purified by adsorption to Speed-amide SPE columns and detected by normal-phase high-performance liquid chromatography, as described by Caputo *et al* [36].

The amount of glucosylation was measured in comparison to control by measuring the peak area of the PNGase F released 2-AA-labelled species Man_9_GlcNAc_2_ and Glc_1_Man_9_GlcNAc_2_ using Waters Empower software. This allows the % of glucosylation to be determined as the % of Glc_1_-species (Peak Area Glc_1_Man_9_GlcNAc_2_) as a total of potential glucosylation species (Peak Area of Glc_1_Man_9_GlcNAc_2_ + Man_9_GlcNAc_2_).

### X-ray Diffraction Data Collection and processing

#### *Ct*UGGT_Kif_

diffraction data were collected on I04@DLS, at a wavelength λ=0.9763 Å, beam size 80×20µm, 0.2° oscillation. Batches 2,3: plate set at 2.9 Å max resolution; batches 4,5, plate set at 3.5 Å max resolution. Batch 2: 450 images, 0.1 s exposure, Transmission T=70%. Batches 3,4: 500 images, 0.2 s exposure, T=100%. Batch 5: 350 images, 0.5 s exposure, T=100%. Recentring followed after each exposure.

#### *Ct*UGGT-ΔTRXL2

data were collected on I04@DLS, at a wavelength λ=0.97950 Å, beam size 43×30 µm, 0.15° oscillation, 1200 images, 0.02 s/image and T=100%; plate set at 4.5 Å max resolution.

#### *Ct*UGGT^S180C/T742C^

data were collected on I24@DLS, at a wavelength λ=0.96860 Å, beam size 50×50 µm, 0.10° oscillation, 1800 images, 0.1 s/image and T=30%; plate set at 3.5 Å max resolution.

#### *Ct*UGGT^G177C/A786C^

data were collected on I04@DLS, at wavelength λ=0.97949 Å, beam size 19×10 µm, 0.10° oscillation, 1800 images, 0.10 s/image and T=100%; plate set at 4.5 Å max resolution.

All datasets were processed with the `autoPROC` suite of programs [37]. SI Appendix Table S1 contains the data processing statistics.

### Crystal Structure Determination and Refinement

#### *Ct*UGGT_Kif_

(PDB ID 6TRF): `Phaser` [38] was run in all primitive orthorhombic space groups searching for one copy of PDB ID 5NV4 from which TRXL2 was removed (declaring a RMSD_C_α of 2.0 Å - Phaser refined it to 0.77 Å). The results were clearly best in P2_1_2_1_2_1_ (RF Z-score 7.0; TF Z-score: 10.4; Refined TFZ-equiv: 16.3; LLG: 114; Refined TF Z-score: 16.3, Refined LLG: 208. wR=0.58). The first map obtained in `autoBUSTER` from this MR model (which lacks TRXL2) showed strong density for the TRXL2 domain. The TRXL2 domain was added by superposing PDB ID 5NV4 onto the model, and real-space fitting the domain to the Fo-Fc map in `CCP4-coot` [45]. The structure was refined in `autoBUSTER` [39] with one TLS body per domain with external restraints [40] to PDB ID 5NV4.

#### *Ct*UGGT-ΔTRXL2

(PDB ID 6TS2): Molecular replacement with the program `CCP4-Molrep` was initially attempted using *Ct*UGGT PDB entry 5NV4 with the TRXL2 domain removed. Electron density for the TRXL3 domain (residues 667-879) was poor. This suggested that upon deletion of TRXL2, the relative orientation of TRXL3 with respect to the rest of the protein was also changed. The TRXL3 domain was therefore also cut from the search model. Three copies of this model were placed with `CCP4-Molrep`. A first round of refinement was carried out in `autoBUSTER` with one TLS body per domain, and one rigid body per domain, with automated NCS restraints and external secondary structure restraints to the deposited 5NV4 structure (R=35.0%, Rfree=37.6%). The phases showed positive difference density in regions close to the loose ends of the search model on either side of TRXL3 for copies A,B,C, suggesting that indeed the deletion of TRXL2 caused TRXL3 to rearrange. Two copies of the TRXL3 domain were then placed with `CCP4-Molrep`, clearly belonging to two of the molecules in the asymmetric unit. An additional search for a third TRXL3 copy gave a convincing solution that did not appear to belong to the three molecules so far placed, highlighting the possible presence of a fourth copy in the asymmetric unit. This model comprising two copies of *Ct*UGGT-ΔTRXL2, a *Ct*UGGT-ΔTRXL2-ΔTRXL3 model and a TRXL3 domain for a fourth copy was subject to refinement with the same protocol as above (R=31.9% Rfree=33.3%). After this refinement, electron density for the missing TRXL3 domain and the remaining domains of the fourth copy of the molecule was visible in the map. One of the *Ct*UGGT-ΔTRXL2 molecules was superposed onto the fourth copy’s TRXL3 domain, followed by rigid body fitting of the bulk of the final copy in `Coot` [45]. The final model was refined in `autoBUSTER` with one set of TLS thermal motion tensors per domain and non-crystallographic symmetry and external restraints to the PDB ID 5NV4 structure.

#### *Ct*UGGT^S180C/T742C^

(PDB ID 6TRT): `CCP4-Molrep` was run against the *Ct*UGGT^S180C/T742C^ data in P3_1_12 and P3_2_12 searching with a copy of PDB ID 5NV4 from which all three TRXL1,2,3 were removed (leaving only TRXL4, BS1,BS2 and GT24 domains). The results were clearly better in P3_2_12 (P3_2_12 has wR=0.606, Score=0.435, TF/sigma=10.35, Contrast=9.21 *vs.* P3_1_12 wR=0.637, Score=0.372, TF/sigma=5.61, Contrast=4.76). The first electron density map obtained in `autoBUSTER` [41] from this MR model shows strong density for the TRXL3 domain and the Ca^2+^ site in the GT24 domain. The TRXL3 domain was added by superposing PDB ID 5NV4 onto the model, and real-space fitting the domain to the Fo-Fc map in `CCP4-coot` [45]. After one more round of refinement, the TRXL1 domain was added in the same way. Finally, the TRXL2 domain was added by molecular replacement with `CCP4-Molrep` in presence of the rest of the structure. The structure was refined in `autoBUSTER` [41] with one TLS body per domain, one rigid body per domain, with external restraints [40] to PDB ID 5NV4. Fo-Fc residuals on two sites (the catalytic site and a crystal contact between TRXL2 and one of its symmetry mates) suggested a lanthanide ion from the crystallisation mix (which contains Y^3+^, Tb^3+^, Er^3+^, Yb^3+^). The ions are likely either Er^3+^ or Tb^3+^, which are known to substitute for Ca^2+^ and Mn^2+^ in protein coordination sites [42–44]. At the wavelength of data collection, λ=0.96861 Å, f’_Er3+_=-1.7235 e^-^ and f’’_Er3+_=8.2682 e^-^, while f’_Tb3+_=-1.046 e^-^ and f’’_Tb3+_=6.9753 e^-^. Peaks at +9.4 and +7.4 sigmas are indeed visible at these two sites in the anomalous Fourier difference map. The ions were modelled as Tb^3+^, with a Tb^3+^-O distance of 2.4 ± 0.3 Å (coordinating residues: site 1: D1302, D1304, D1435; site 2: E774 from a symmetry mate, E713, E716 and D818).

#### *Ct*UGGT^G177C/A786C^

(PDB ID): The *Ct*UGGT^G177C/A786C^ crystal structure was initially phased by Molecular Replacement with `Molrep` searching in space group P4_3_2_1_2 for one copy of PDB ID 5NV4 from which TRXL1 and TRXL2 were removed. The first map obtained in `autoBUSTER` [41] from this MR model showed density for the missing domains, which were added by superposing PDB ID 5NV4 onto the model, and real-space fitting the TRXL1 and TRXL2 domains to the Fo-Fc map in `CCP4-coot` [45]. The structure was refined in `autoBUSTER` [39] with one TLS body per domain with local automated NCS restraints and external restraints [40] to PDB ID 5NV4. Portions of the catalytic domain are disordered in the crystal and could not be traced.

SI Appendix Table S2 reports the Rfactors and geometry statistics for all models after the final refinements.

### Fitting of the *Td*UGGT structure in the negative stain EM map

The crystal structures of *Td*UGGT catalytic domain (PDB ID 5H18, residues 1190-1466) and of the *Td*UGGT N-terminal portion (PDB ID 5Y7O, residues 29-1042) were aligned with the full-length *Ct*UGGT intermediate structure (PDB ID 5MU1, residues 1190-1466) in `Coot` [45]. `Modeller` [46] was then used to complete the *Td*UGGT structure, homology modelling the missing portions 158-165; 251-282; 403-414; 684-693; 738-741; 756-759; 1038-1150; 1380-1384. The fittings of the *Td*UGGT and anti-*Td*UGGT Fab models into the negative stain EM map and in its inverse hand were carried out in `Chimera` [47]. Both for original and inverse hand map, the *Td*UGGT homology model was first aligned manually with the map, low-pass filtered to a resolution of 25 Å, then fitted to the EM map using the `Fit in map` tool in `Chimera`. After fitting the TdUGGT model, a Fab model from PDB ID 1FGN was fitted to the map with the same `Fit in map` tool in `Chimera`, again after low-pass filtering the PDB model to 25 Å. Final real space CCs in the original and inverse hand maps: ^ori^CC*_Td_*_UGGT_=0.89; ^ori^CC_Fab_=0.90; ^inv^CC*_Td_*_UGGT_=0.90; ^inv^CC_Fab_=0.90.

### Computational Simulations

#### System Preparation

We used as initial structures the four available *Ct*UGGT structures [5], which we call ‘open’ (PDB ID 5MZO); ‘intermediate’ (PDB ID 5MU1); ‘closed’ (PDB ID 5N2J); the mutant D611C-G1050C ‘closed-like’ (PDB ID 5NV4); and the newly determined ‘new intermediate’ *Ct*UGGT_Kif_ (PDB ID 6TRF). Starting from each structure, we performed 250 ns all-atom Molecular Dynamics (MD) simulations using the AMBER force field and package [48] and analyzed the resulting dynamics and conformational landscape using Principal Component (PC) analysis. For each system, all non-protein molecules (carbohydrates and ions) were removed from the crystal structure. Gap regions (residues 242-273, 1152-1187 and 1334-1342) were completed and refined using the Modeller v9.19 software [49]. Standard protonation states were assigned to titratable residues (Asp and Glu are negatively charged; Lys and Arg are positively charged). Histidine protonation was assigned favoring formation of hydrogen bonds in the crystal structure. The complete protonated systems were then solvated by a truncated cubic box of TIP3P waters, ensuring that the distance between the biomolecule surface and the box limit was at least 10 Å.

#### MD simulations

Each system was first optimized using a conjugate gradient algorithm for 5000 steps, followed by 150 ps. long constant volume MD equilibration, in which the first 100 ps were used to gradually raise the temperature of the system from 0 to 300 K (integration step = 0.0005 ps/step). The heating was followed by a 250 ps. long constant temperature and constant pressure MD simulation to equilibrate the system density (integration step = 0.001 ps/step). During these temperature and density equilibration processes, the protein alpha-carbon atoms were constrained by 5 kcal/mol/Å force constant using a harmonic potential centered at each atom starting position. Next, a second equilibration MD of 500 ps. was performed, in which the integration step was increased to 2 fs and the force constant for restrained alpha-carbons was decreased to 2 kcal/mol/Å. Finally, a 1 ns. long MD simulation was carried out with no constraints and the ‘Hydrogen Mass Repartition’ technique, which allows an integration step of 4 fs, and these settings were kept for all the subsequent Production 20 ns long MD runs.

All simulations were performed with the Amber package of programs using the ff14SB force field for all amino acid residues. Pressure and temperature were kept constant using the Monte-Carlo barostat and Langevin thermostat, respectively, using the Amber default coupling parameters. All simulations were performed with a 10 Å cutoff for nonbonded interactions, and periodic boundary conditions using the Particle Mesh Ewald summation method for long-range electrostatic interactions. The SHAKE algorithm was applied to all hydrogen-containing bonds in all simulations with an integration step equal or higher than 2 fs.

#### PC calculations

All trajectory processing and PC calculations were performed with the Cpptraj (Roe and Cheatham, 2013) module of the AMBER package. For each individual MD, PCs of the alpha-carbons were computed over an ensemble of 6000 trajectory frames representing the 250 ns long trajectories.

### Mass spectroscopy: tryptic peptides

Protein samples were digested in-solution with sequencing grade trypsin (Promega). Briefly: samples were treated in 100 mM iodo-acetamide for 1 hour in dark to alkylate any free cysteines followed by denaturing with 8M urea for 40 min. The samples were further diluted with 50mM ammonium bicarbonate to reduce the Urea concentration to 1M. 1uL of 300ng/uL trypsin solution was added to each sample and incubated at 37 °C overnight. The resulting samples were directly analysed by LC-MS.

Tryptic peptides of *Ct*UGGT, and the double mutants *Ct*UGGT^G177C/A786C^, *Ct*UGGT^G179C/T742C^ and *Ct*UGGT^S180C/T742C^ were separately run on an Dionex UltiMate3000 RSLC (Thermo Scientific) and electrosprayed directly into a Q Exactive mass spectrometer (Thermo Fischer Scientific) through an Flex ion-electrospray ion source (Thermo Fischer Scientific). Peptides were trapped on a C18 PepMap trapping column (µ-Precolumn, 300 µM I.D. x 5 mm, 100 µm particle size, 100 Å, Thermo Scientific) at a flow rate 10 µL/min. The trapping buffer was 0.05% v/v trifluoroacetic acid (TFA) in water (LC-MS grade). Samples were then separated using a C18, 75 µm x 25 cm (Acclaim PepMap nanoViper, part number 164941, 2.0 µm particle size, 100 Å, Thermo Scietific) analytical column (with mobile phases: 0.1% Formic acid in water (A) and 0.1% formic acid in acetonitrile (B)) at a flow rate of 300 nL/min, and the following gradients: minutes (mins) 0-5.5: 2% B; mins 5.5-10: 8% B; mins 10-40: 45% B; mins 40-41: 95% B; mins 41-46: 95% B; mins 46-60: 2% B.

Data were acquired in Data Dependent Mode (DDA) using the following settings: chromatographic peak width: 20 s; resolution: 70,000; AGC target: 3×10^6^; maximum IT (injection time): 100 ms; scan range: 300 to 2000 m/z; ddMS2 resolution: 17,500; AGC target: 5×10^4^; maximum IT: 100 ms; loop count: 10 (*i.e.* Top 10); isolation width: 4.0 m/z; fixed first mass: 120.0 m/z. Data dependent (dd) settings: minimum AGC target: 5.0×10^3^; intensity threshold: 5.0×10^4^; charge exclusion: 1; peptide match: preferred; exclude isotope: on; dynamic exclusion: 30.0 s. A normalized Collision energy (NCE) of 27 was used for the fragmentation of peptides in a high-energy collision dissociation (HCD) cell and the s-lens setting in the tune file was changed to 70.

#### Data analysis (crosslinking and protein identification)

MassMatrix (version 2.4.2) was used for data analysis to find S-S cross linking and protein/peptide identification. A customized database, containing the sequences of the proteins of interest, was used to perform searches. MS data were converted into .mgf format using MSconvert from the ProteoWizard toolbox [50]. Search parameters were as follows: maximum number of missed cleavages = 4; fixed modification = none; variable modifications: CAMC-Iodoacetamide derivative (Carbamidomethyl) of C and OxiM-Oxidation of M; disulphide bonds were considered as the crosslink (Cys-Cys, −2.02 Da); mass accuracy filter = 20 ppm for precursor ions; MS2 tolerance = 0.02 Da (values as per the Massmatrix user’s protocol). The quality of a peptide match is mainly evaluated by three statistical scores: pp, pp2, pptag. A peptide match with max (pp,pp2) > 2.7 and pptag > 1.3 is considered to be significant with p value < 0.05.

## Supporting information

SI Appendix movie III

## Acknowledgments

We thank Julio J. Caramelo, Alessandro T. Caputo and Raymond Dwek, for helpful discussions, advice and comments on the manuscript and the members of the Zitzmann laboratory for assistance with molecular biology and protein chemistry. Yentli Soto Albrecht designed the *Ct*UGGT full-length WT sequencing primers. Ed Lowe and the staff at beamlines I03, I04 and I24 at the Diamond Light Source, Harwell, England, UK assisted with X-ray data collection. Tadashi Satoh and Koichi Kato kindly provided us with their 25 Å negative stain EM reconstruction of the complex between *Td*UGGT and its Fab. P.R. is the recipient of a Leicester LISCB-Wellcome Trust ISSF award, grant reference 204801/Z/16/Z. G.T. is funded by the 18-months Wellcome Trust Seed Award in Science 214090/Z/18/Z to P.R. J.C.H. was funded by the Wellcome Trust 4-year Studentship 106272/Z/14/Z. N.Z. is a Fellow of Merton College, Oxford.

## Author Contributions

P.R., M.A.M., A.S. and N.Z. conceived and designed the study. M.A.M, C.P.M. and J.I.B.C. carried out the molecular dynamics experiments. L.M., P.R., S.R., A.L. and J.R. cloned the *Ct*UGGT-ΔTRXL2 construct. P.R., S.R., A.L., S.V. and J.R. cloned, expressed, purified and determined the crystal structure of *Ct*UGGT-ΔTRXL2. P.R., A.L., J.C.H., J.I.B.C. and G.T. fitted the *Td*UGGT:Fab structure in the *Td*UGGT:Fab negative-stained EM map. P.R., S.V., L.M., A.L., R.I., A.V.C and M.H. cloned, expressed, purified and determined the crystal structures of the *Ct*UGGT^G177C/A786C^ and *Ct*UGGT^S180C/T742C^ mutants. A.K. collected and analysed the mass spectra of the double Cys *Ct*UGGT mutants. P.R., J.R., R.I., A.L. and D.S.A carried out re-glucosylation assays. All authors contributed to the writing of the paper.

---

**SI Appendix Figure S1.**
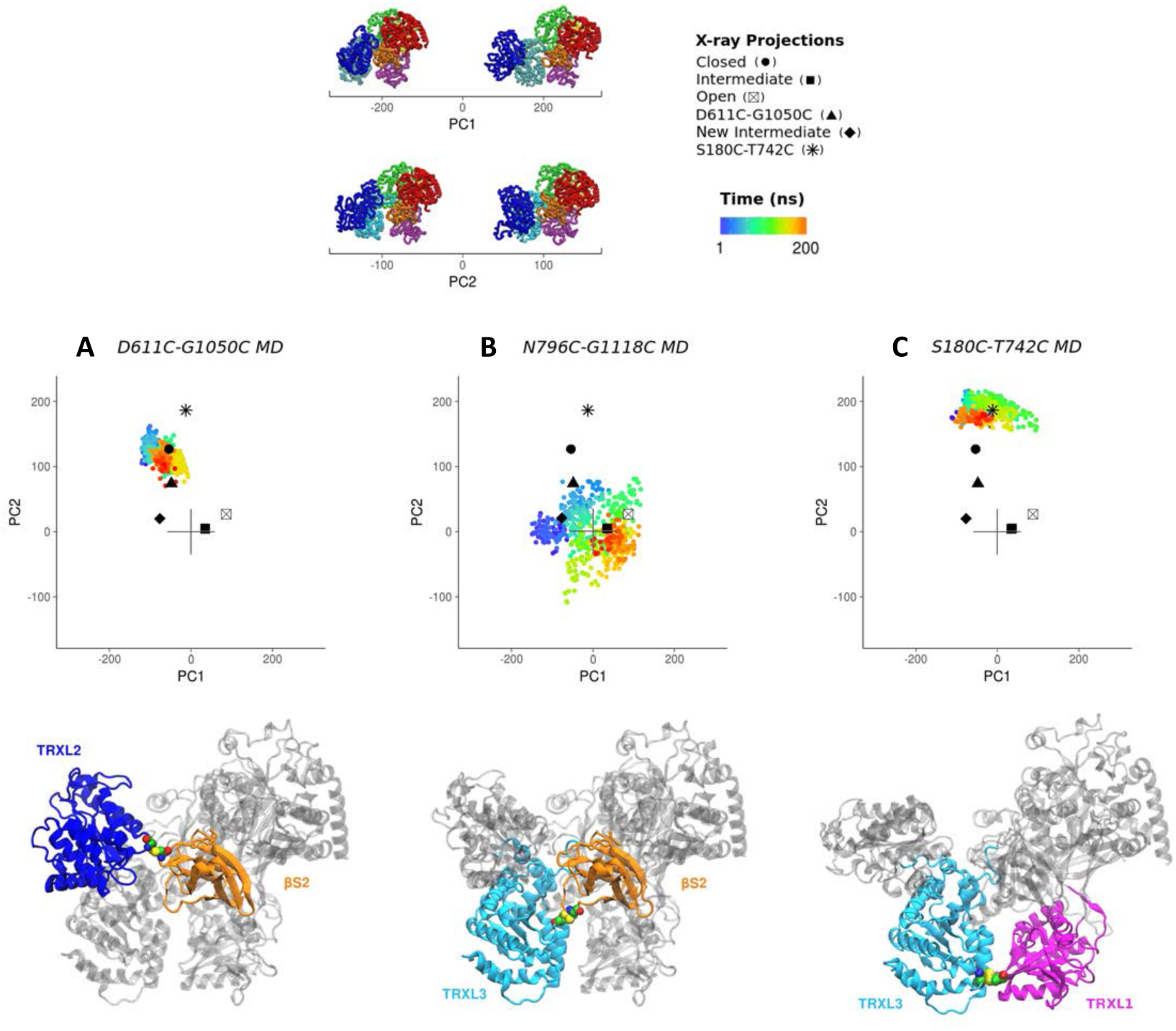
Projections of individual MD trajectories for *Ct*UGGT double Cysteine mutants onto the full conformational landscape of the wild type enzyme, coloured as a function of time. Lower panels show the structure of each mutant, with the mutated cysteine residues drawn in sphere representation and domains containing the mutation shown in colour, with the rest of the protein is in grey. **(A)** Crystal structure of the *Ct*UGGT^D611C/G1050C^ mutant (PDB ID: 5NV4). **(B)** Model structure of the *Ct*UGGT^N796C-G1118C^ mutant, generated using the closed X-ray structure as template and the Modeller v.9.19 Software. **(C)** Crystal structure of the *Ct*UGGT^S180C/T742C^ mutant (PDB ID: 6TRT).

**SI Appendix Figure S2.**
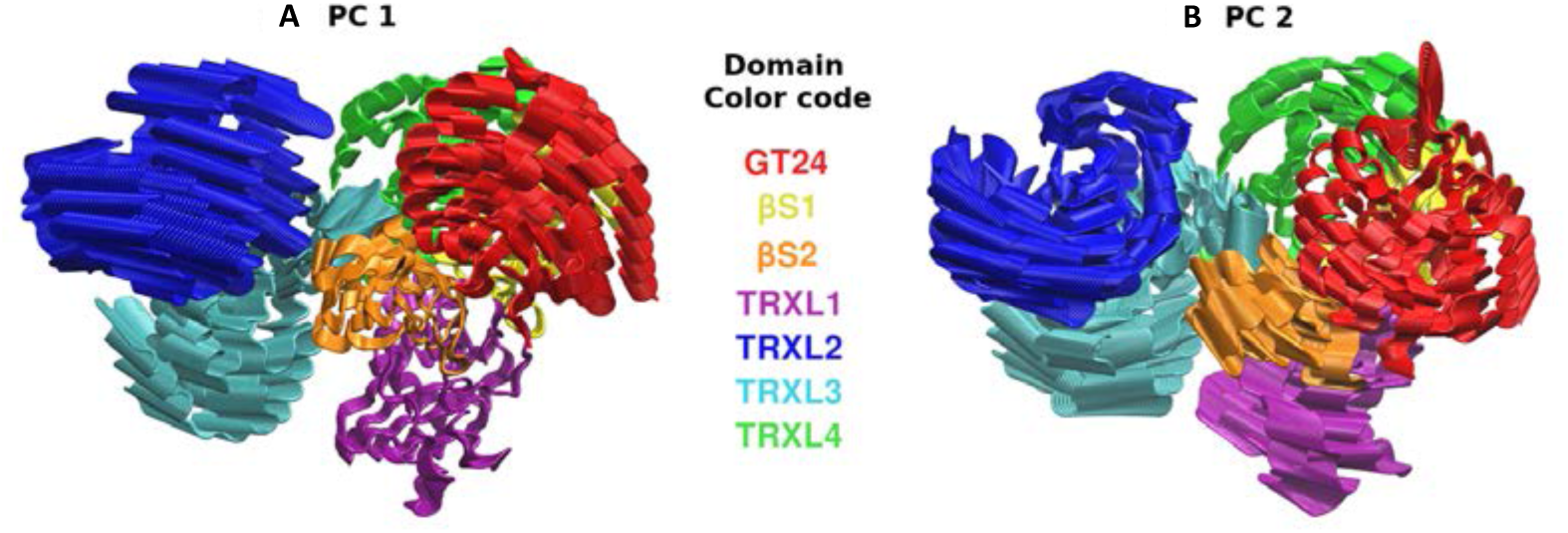
The first two principal components (PCs) of the joint MDs. Domains coloured as in Figure 1A.

**SI Appendix FIgure S3.**
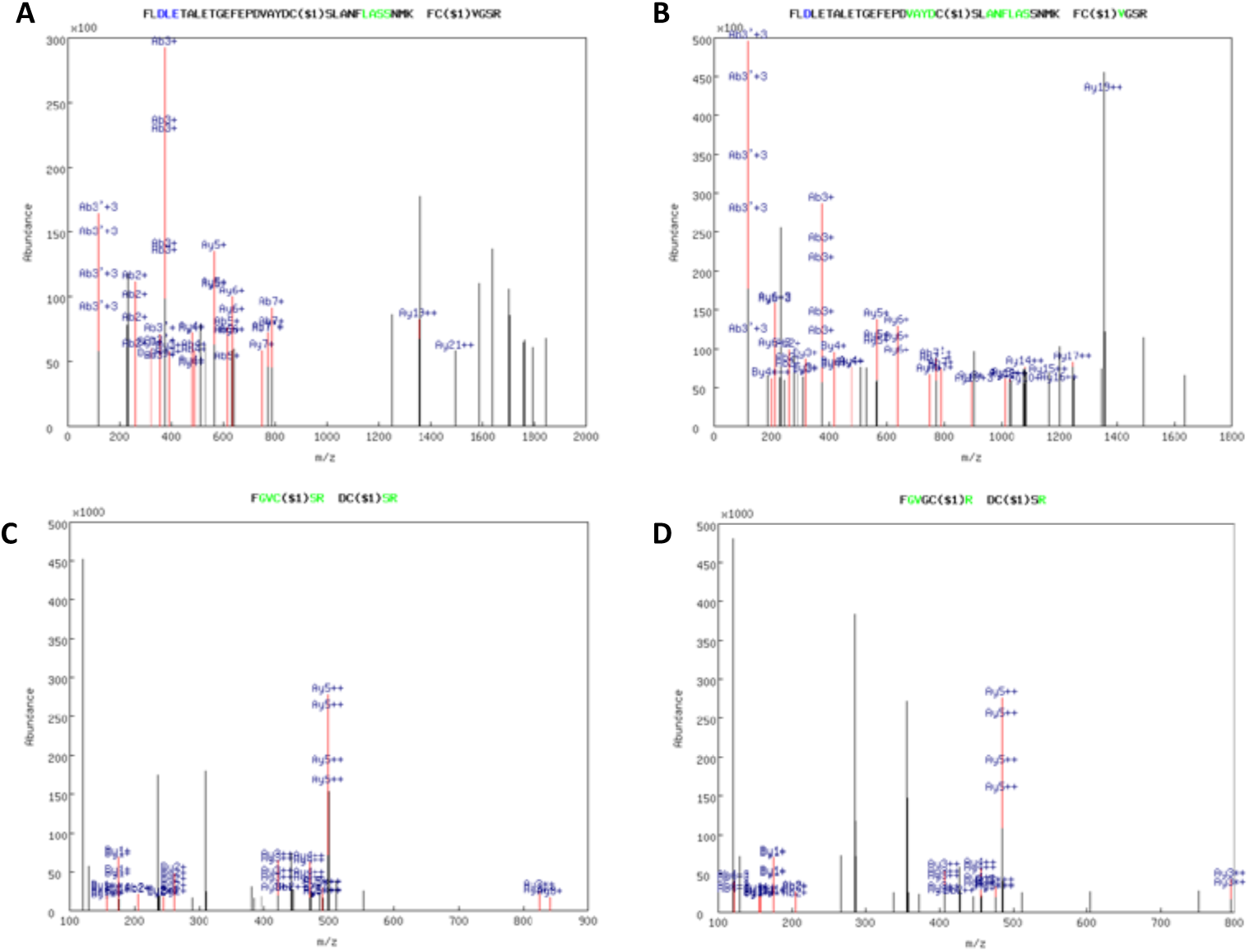
Mass spectrometry of tryptic peptides confirms the disulfides in the *Ct*UGGT double Cys mutants *Ct*UGGT^G177C/A786C^, *Ct*UGGT^G179C/T742C^ and *Ct*UGGT^S180C/T742C^. In peptide mass spectrometry, fragment ions that appear to extend from the amino- or carboxy-terminus of a peptide are termed “b” or “y” ions, respectively. (A,B) Mass spectrometry detection of ions derived from fragmentation of the disulphide-bridged tryptic peptides ^766^FLDLETALETGEFEPDVAYDCSLANFLASSNMK^798^ and ^176^FCVGSR^181^ in the double mutant *Ct*UGGT^G177C/A786C^. The ions confirm the establishment of the engineered disulphide bridge at positions 177-786 between the TRXL1 and TRXL3 domains. No peptides containing free Cys at either position 177 or 786 were detected. (C) Mass spectrometry detection of ions derived from fragmentation of the disulphide-bridged tryptic peptides ^741^DCSR^744^ and ^176^FGVCSR^181^ in the double mutant *Ct*UGGT^G179C/T742C^. The ions confirm the establishment of the engineered disulphide bridge at positions 179-742 between the TRXL1 and TRXL3 domains. No peptides containing free Cys at either position 179 or 742 were detected. (D) Mass spectrometry detection of ions derived from fragmentation of the disulphide-bridged tryptic peptides ^741^DCSR^744^ and ^176^FGVGCRDVILYADITS^191^ in the double mutant *Ct*UGGT^S180C/T742C^. The ions confirm the establishment of the engineered disulphide bridge at positions 180-742 between the TRXL1 and TRXL3 domains. No peptides containing free Cys at either position 180 or 742 were detected.

**SI Appendix Figure S4.**
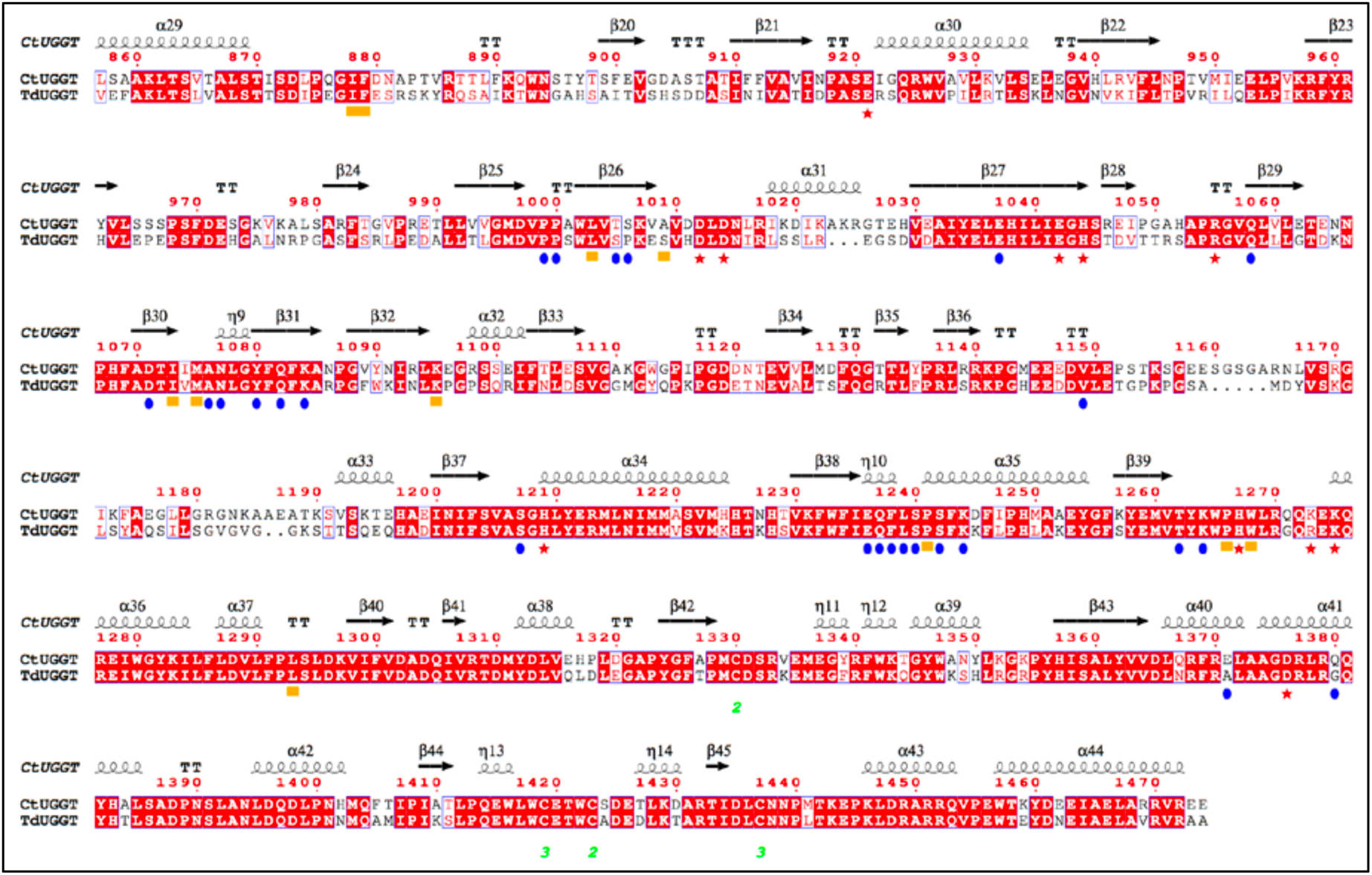
Sequence conservation at the interface of the GT24 and βS1-βS2 domains. The C-terminal parts of the sequences of *Ct*UGGT and *Td*UGGT (centred around the residues in the GT24:βS1-βS2 interface) are aligned and the conserved residues shown in white text over red squares. Similar residues are in red text over white squares with blue edges. The red numbers indicate *Ct*UGGT sequence numbers. Disulphide bonds are labelled in green under the Cys residues. The *Ct*UGGT secondary structure is indicated above its sequence. Blue dots: residues whose side chains are forming hydrogen bonds across the GT24:βS1-βS2 domains interface. Red stars: residues whose side chains are forming salt bridges across the GT24:βS1-βS2 domains interface. Orange squares: residues whose side chains are forming hydrophobic interactions across the GT24:βS1-βS2 domains interface. The sequences were aligned using `Clustal Omega` [52]. The figure has been made using `ESPript` [53].

## SI Appendix Tables

**SI Appendix Table S1.** *Ct*UGGT X-ray diffraction data collection statistics.

| CtUGGT Structure | CtUGGT<sub>Kif</sub> | ΔTRXL2 | G177C/A786C | S180C/T742C |
| --- | --- | --- | --- | --- |
| PDB ID | 6TRF | 6TS2 | 6TS8 | 6TRT |
| Beamline, date | I03@DLS,<br>01.05.2016 | I04@DLS,<br>13.01.2018 | I04@DLS,<br>08.10.2018 | I24@DLS,<br>08.08.2018 |
| Space group (Z) | P2<sub>1</sub>2<sub>1</sub>2<sub>1</sub> (4) | P2<sub>1</sub> (8) | P4<sub>3</sub>2<sub>1</sub>2 (8) | P3<sub>2</sub>12 (6) |
| Wavelength (Å) | 0.97630 | 0.97950 | 0.97949 | 0.96861 |
| Cell dimensions<br>a, b, c (Å) | a=78.65,<br>b=148.93,<br>c=190.30 | a=151.14<br>b=191.01<br>c=158.81 | a=b=138.22<br>c=175.44 | a=b=148.80<br>c=235.55 |
| $\alpha$, $\beta$, $\gamma$ (°) | $\alpha$=90.0,<br>$\beta$=90.0, $\gamma$=90.0 | $\alpha$=90.0, $\beta$=117.,<br>$\gamma$=90.0 | $\alpha$=90.0,<br>$\beta$=90.0, $\gamma$=90.0 | $\alpha$=90.0,<br>$\beta$=90.0, $\gamma$=120.0 |
| Resolution range (Å) | 95.15-4.11 (4.49-4.11) | 140.61-5.74 (6.15-5.74) | 108.87-4.46 (5.13-4.46) | 128.86-4.58 (5.13-4.58) |
| $R_{\text{merge}}$ | 0.149 (3.268) | 0.150 (1.432) | 0.314 (1.474) | 0.118 (1.563) |
| $R_{\text{meas}}$ | 0.157 (3.376) | 0.189 (2.146) | 0.337 (1.546) | 0.125 (1.650) |
| $CC_{1/2}$ | 0.997 (0.566) | 0.994 (0.382) | 0.996 (0.774) | 0.995 (0.411) |
| $I / \sigma(I)$ | 9.9 (1.3) | 3.6 (1.3) | 5.1 (1.6) | 10.6 (1.6) |
| Completeness (%) | 90.2 (74.2) | 91.5 (52.6) | 90.2 (92.5) | 90.1 (83.0) |
| Redundancy | 10.9 (15.9) | 5.6 (6.0) | 7.7 (11.1) | 8.9 (9.8) |

**SI Appendix Table S2.** *Ct*UGGT crystal structures, refinement statistics.

| **Crystal form** | **CtUGGT<sub>Kif</sub>** | **ΔTRXL2** | **G177C/A786C** | **S180C/T742C** |
| --- | --- | --- | --- | --- |
| **PDB ID** | 6TRF | 6TS2 | 6TS8 | 6TRT |
| **Space group (Z)** | P2<sub>1</sub>2<sub>1</sub>2<sub>1</sub> (4) | P2<sub>1</sub> (8) | P4<sub>3</sub>2<sub>1</sub>2 (8) | P3<sub>2</sub>12 (6) |
| **Resolution (Å)** | 95.1-4.1<br>(4.6-4.1) | 140.6-5.7<br>(6.0-5.7) | 108.4-4.5<br>(5.6-4.5) | 128.9-4.6<br>(5.1-4.6) |
| **No. reflections** | 7,503 (442) | 17,358 (424) | 3,573 (397) | 9,528 (477) |
| **R<sub>work</sub> / R<sub>free</sub>** | 0.25/0.31<br>(0.26/0.22) | 0.17/0.24<br>(0.22/0.33) | 0.28/0.29<br>(0.29/0.40) | 0.29/0.30<br>(0.25/0.26) |
| **Atoms** | 10,717 | 35,718 | 10,478 | 11,210 |
| **B-factors (Å<sup>2</sup>)** | 270 | 134 | 119 | 143 |
| **Rmsd<sub>bonds</sub> (Å)** | 0.006 | 0.009 | 0.02 | 0.006 |
| **Rmsd<sub>angles</sub> (°)** | 0.95 | 1.07 | 1.76 | 0.99 |
| **Ramachandran outliers** | 17/1309<br>(1.3%) | 87/4431<br>(1.96%) | 15/1285<br>(1.2%) | 14/1363<br>(1.0%) |
| **Ramachandran allowed** | 1292/1309<br>(98.7%) | 4344/4431<br>(98.0%) | 1270/1285<br>98.8% | 1349/1363<br>(99.0%) |
| **Ramachandran favoured** | 1235/1309<br>(94.3%) | 3934/4431<br>(88.8%) | 1196/1285<br>(93.1%) | 1284/1363<br>(94.2%) |

All structures were refined against X-ray data from one crystal only. Values in parentheses are for highest-resolution shell.

**SI Appendix Table S3.**
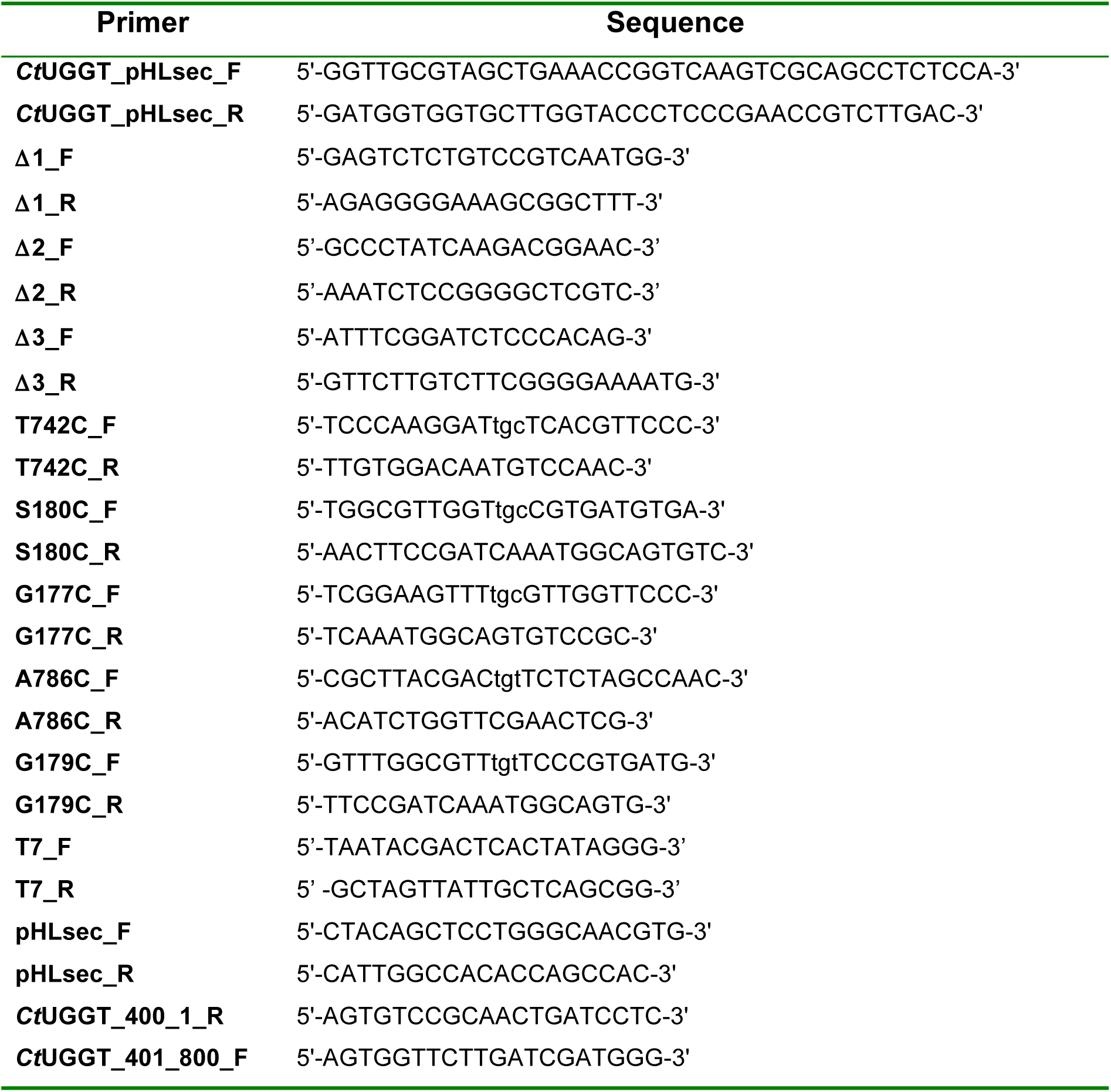
DNA primers used in this work.

## References

1. Vincenz-Donnelly L, Hipp MS (2017) The endoplasmic reticulum: A hub of protein quality control in health and disease. Free Radic Biol Med 108:383–393. doi: 10.1016/j.freeradbiomed.2017.03.031

2. Amara JF, Cheng SH, Smith AE (1992) Intracellular protein trafficking defects in human disease. Trends Cell Biol 2:145–149.

3. Parodi AJ, Caramelo JJ, D’Alessio C (2014) UDP-Glucose: Glycoprotein Glucosyltransferase 1,2 (UGGT1,2). In: Handbook of Glycosyltransferases and Related Genes. Springer Japan, Tokyo, pp 15–30

4. Calles-Garcia D, Yang M, Soya N, et al (2017) Single-particle electron microscopy structure of UDP-glucose:glycoprotein glucosyltransferase suggests a selectivity mechanism for misfolded proteins. J Biol Chem jbc.M117.789495. doi: 10.1074/jbc.M117.789495

5. Roversi P, Marti L, Caputo AT, et al (2017) Interdomain conformational flexibility underpins the activity of UGGT, the eukaryotic glycoprotein secretion checkpoint. Proc Natl Acad Sci USA. doi: 10.1073/pnas.1703682114

6. Satoh T, Song C, Zhu T, et al (2017) Visualisation of a flexible modular structure of the ER folding-sensor enzyme UGGT. Sci Rep 7:12142. doi: 10.1038/s41598-017-12283-w

7. Satoh T, Kato K (2018) Structural Aspects of ER Glycoprotein Quality-Control System Mediated by Glucose Tagging. Adv Exp Med Biol 1104:149–169. doi: 10.1007/978-981-13-2158-0_8

8. Capece L, Estrin DA, Marti MA (2008) Dynamical characterization of the heme NO oxygen binding (HNOX) domain. Insight into soluble guanylate cyclase allosteric transition. Biochemistry 47:9416–9427. doi: 10.1021/bi800682k

9. Krissinel E (2015) Stock-based detection of protein oligomeric states in jsPISA. Nucleic Acids Res 43:W314–9. doi: 10.1093/nar/gkv314

10. Rosenthal PB, Henderson R (2003) Optimal determination of particle orientation, absolute hand, and contrast loss in single-particle electron cryomicroscopy. Journal of Molecular Biology 333:721–745. doi: 10.1016/j.jmb.2003.07.013

11. Trombetta SE, Bosch M, Parodi AJ (1989) Glucosylation of glycoproteins by mammalian, plant, fungal, and trypanosomatid protozoa microsomal membranes. Biochemistry 28:8108–8116.

12. Parodi AJ (2007) How I became a biochemist. TBMB 59:361–363. doi: 10.1080/15216540601080198

13. Taylor SC, Ferguson AD, Bergeron JJM, Thomas DY (2004) The ER protein folding sensor UDP-glucose glycoprotein–glucosyltransferase modifies substrates distant to local changes in glycoprotein conformation. Nat Struct Mol Biol 11:128–134. doi: 10.1038/nsmb715

14. Ritter C, Helenius A (2000) Recognition of local glycoprotein misfolding by the ER folding sensor UDP-glucose:glycoprotein glucosyltransferase. Nat Struct Biol 7:278–280. doi: 10.1038/74035

15. Ritter C, Quirin K, Kowarik M, Helenius A (2005) Minor folding defects trigger local modification of glycoproteins by the ER folding sensor GT. The EMBO Journal 24:1730–1738. doi: 10.1038/sj.emboj.7600645

16. Totani K, Ihara Y, Matsuo I, Ito Y (2006) Substrate specificity analysis of endoplasmic reticulum glucosidase II using synthetic high mannose-type glycans. Journal of Biological Chemistry 281:31502–31508. doi: 10.1074/jbc.M605457200

17. Totani K, Ihara Y, Tsujimoto T, et al (2009) The Recognition Motif of the Glycoprotein-Folding Sensor Enzyme UDP-Glc:Glycoprotein Glucosyltransferase †. Biochemistry 48:2933–2940. doi: 10.1021/bi8020586

18. Izumi M, Komaki S, Okamoto R, et al (2016) Synthesis of misfolded glycoprotein dimers through native chemical ligation of a dimeric peptide thioester. Org Biomol Chem 14:6088–6094. doi: 10.1039/c6ob00928j

19. Izumi M, Oka Y, Okamoto R, et al (2016) Synthesis of Glc1Man9-Glycoprotein Probes by a Misfolding/Enzymatic Glucosylation/Misfolding Sequence. Angew Chem Int Ed Engl 55:3968– 3971. doi: 10.1002/anie.201511491

20. Kiuchi T, Izumi M, Mukogawa Y, et al (2018) Monitoring of Glycoprotein Quality Control System with a Series of Chemically Synthesized Homogeneous Native and Misfolded Glycoproteins. J Am Chem Soc 140:17499–17507. doi: 10.1021/jacs.8b08653

21. Takeda Y, Seko A, Fujikawa K, et al (2016) Effects of domain composition on catalytic activity of human UDP-glucose:glycoprotein glucosyltransferases. Glycobiology 26:999–1006. doi: 10.1093/glycob/cww069

22. Tax G, Lia A, Santino A, Roversi P (2019) Modulation of ERQC and ERAD: A Broad-Spectrum Spanner in the Works of Cancer Cells? J Oncol 2019:8384913. doi: 10.1155/2019/8384913

23. Keith N, Parodi AJ, Caramelo JJ (2005) Glycoprotein tertiary and quaternary structures are monitored by the same quality control mechanism. Journal of Biological Chemistry 280:18138–18141. doi: 10.1074/jbc.M501710200

24. Zhang W, Wearsch PA, Zhu Y, et al (2011) A role for UDP-glucose glycoprotein glucosyltransferase in expression and quality control of MHC class I molecules. Proc Natl Acad Sci USA 108:4956–4961. doi: 10.1073/pnas.1102527108

25. Gardner TG, Kearse KP (1999) Modification of the T cell antigen receptor (TCR) complex by UDP-glucose:glycoprotein glucosyltransferase. TCR folding is finalized convergent with formation of alpha beta delta epsilon gamma epsilon complexes. Journal of Biological Chemistry 274:14094–14099.

26. Mackeen MM, Almond A, Deschamps M, et al (2009) The conformational properties of the Glc3Man unit suggest conformational biasing within the chaperone-assisted glycoprotein folding pathway. Journal of Molecular Biology 387:335–347. doi: 10.1016/j.jmb.2009.01.043

27. Albesa-Jové D, Guerin ME (2016) The conformational plasticity of glycosyltransferases. Curr Opin Struct Biol 40:23–32. doi: 10.1016/j.sbi.2016.07.007

28. Albesa-Jové D, Sainz-Polo MÁ, Marina A, Guerin ME (2017) Structural Snapshots of α-1,3-Galactosyltransferase with Native Substrates: Insight into the Catalytic Mechanism of Retaining Glycosyltransferases. Angew Chem Int Ed Engl 77:521. doi: 10.1002/anie.201707922

29. Caramelo JJ, Castro OA, Alonso LG, et al (2003) UDP-Glc:glycoprotein glucosyltransferase recognizes structured and solvent accessible hydrophobic patches in molten globule-like folding intermediates. Proceedings of the National Academy of Sciences 100:86–91. doi: 10.1073/pnas.262661199

30. Daikoku S, Seko A, Ito Y, Kanie O (2014) Glycan structure and site of glycosylation in the ER-resident glycoprotein, uridine 5’-diphosphate-glucose: glycoprotein glucosyltransferases 1 from rat, porcine, bovine, and human. Biochemical and Biophysical Research Communications 451:356–360. doi: 10.1016/j.bbrc.2014.07.095

31. Daikoku S, Seko A, Son S-H, et al (2015) The relationship between glycan structures and expression levels of an endoplasmic reticulum-resident glycoprotein, UDP-glucose: Glycoprotein glucosyltransferase 1. Biochemical and Biophysical Research Communications 462:58–63. doi: 10.1016/j.bbrc.2015.04.105

32. Marin MB, Ghenea S, Spiridon LN, et al (2012) Tyrosinase degradation is prevented when EDEM1 lacks the intrinsically disordered region. PLoS ONE 7:e42998. doi: 10.1371/journal.pone.0042998

33. Aricescu AR, Lu W, Jones EY (2006) A time- and cost-efficient system for high-level protein production in mammalian cells. Acta Crystallogr D Biol Crystallogr 62:1243–1250. doi: 10.1107/S0907444906029799

34. Gorrec F (2009) The MORPHEUS protein crystallization screen. J Appl Crystallogr 42:1035–1042. doi: 10.1107/S0021889809042022

35. Gorrec F (2015) The MORPHEUS II protein crystallization screen. Acta Crystallogr F Struct Biol Commun 71:831–837. doi: 10.1107/S2053230X1500967X

36. Caputo AT, Alonzi DS, Marti L, et al (2016) Structures of mammalian ER α-glucosidase II capture the binding modes of broad-spectrum iminosugar antivirals. Proc Natl Acad Sci USA 113:E4630–8. doi: 10.1073/pnas.1604463113

37. Vonrhein C, Flensburg C, Keller P, et al (2011) Data processing and analysis with the autoPROC toolbox. Acta Crystallogr D Biol Crystallogr 67:293–302. doi: 10.1107/S0907444911007773

38. McCoy AJ, Grosse-Kunstleve RW, Adams PD, et al (2007) Phaser crystallographic software. J Appl Crystallogr 40:658–674. doi: 10.1107/S0021889807021206

39. Bricogne G, Blanc E, M B, et al (2017) BUSTER 2.10.3. BUSTER 2.10.3

40. Smart OS, Womack TO, Flensburg C, et al (2012) Exploiting structure similarity in refinement: automated NCS and target-structure restraints in BUSTER. Acta Crystallogr D Biol Crystallogr 68:368–380. doi: 10.1107/S0907444911056058

41. Bricogne G, Blanc E, Brandl M, et al (2011) autoBUSTER. Cambridge United Kingdom. Global Phasing Ltd

42. Colman PM, Weaver LH, Matthews BW (1972) Rare earths as isomorphous calcium replacements for protein crystallography. Biochemical and Biophysical Research Communications 46:1999– 2005.

43. Miake-Lye RC, Doniach S, Hodgson KO (1983) Anomalous x-ray scattering from terbium-labeled parvalbumin in solution. Biophysical Journal 41:287–292. doi: 10.1016/S0006-3495(83)84440-3

44. Hsiao Y-Y, Nakagawa A, Shi Z, et al (2009) Crystal structure of CRN-4: implications for domain function in apoptotic DNA degradation. Mol Cell Biol 29:448–457. doi: 10.1128/MCB.01006-08

45. Emsley P, Lohkamp B, Scott WG, Cowtan K (2010) Features and development of Coot. Acta Crystallogr D Biol Crystallogr 66:486–501. doi: 10.1107/S0907444910007493

46. Webb B, Sali A (2016) Comparative Protein Structure Modeling Using MODELLER. Curr Protoc Protein Sci 86:2.9.1–2.9.37. doi: 10.1002/cpps.20

47. Pettersen EF, Goddard TD, Huang CC, et al (2004) UCSF Chimera--a visualization system for exploratory research and analysis. J Comput Chem 25:1605–1612. doi: 10.1002/jcc.20084

48. Maier JA, Martinez C, Kasavajhala K, et al (2015) ff14SB: Improving the Accuracy of Protein Side Chain and Backbone Parameters from ff99SB. J Chem Theory Comput 11:3696–3713. doi: 10.1021/acs.jctc.5b00255

49. Eswar N, Eramian D, Webb B, et al (2008) Protein structure modeling with MODELLER. Methods Mol Biol 426:145–159. doi: 10.1007/978-1-60327-058-8_8

50. Chambers MC, Maclean B, Burke R, et al (2012) A cross-platform toolkit for mass spectrometry and proteomics. Nature Biotechnology 30:918–920. doi: 10.1038/nbt.2377

51. Harpaz Y, Gerstein M, Chothia C (1994) Volume changes on protein folding. Structure/Folding and Design 2:641–649.

52. Sievers F, Higgins DG (2018) Clustal Omega for making accurate alignments of many protein sequences. Protein Sci 27:135–145. doi: 10.1002/pro.3290

53. Robert X, Gouet P (2014) Deciphering key features in protein structures with the new ENDscript server. Nucleic Acids Res 42:W320–4. doi: 10.1093/nar/gku316

54. Dedola S, Izumi M, Makimura Y, et al (2014) Folding of Synthetic Homogeneous Glycoproteins in the Presence of a Glycoprotein Folding Sensor Enzyme. Angew Chem Int Ed 53:2883–2887. doi: 10.1002/anie.201309665

55. Izumi M, Makimura Y, Dedola S, et al (2012) Chemical synthesis of intentionally misfolded homogeneous glycoprotein: a unique approach for the study of glycoprotein quality control. J Am Chem Soc 134:7238– 7241. doi: 10.1021/ja3013177

56. Pearse BR, Tamura T, Sunryd JC, et al (2010) The role of UDP-Glc:glycoprotein glucosyltransferase 1 in the maturation of an obligate substrate prosaposin. J Cell Biol 189:829–841. doi: 10.1083/jcb.200912105

57. Sousa M, Parodi AJ (1995) The molecular basis for the recognition of misfolded glycoproteins by the UDP-Glc:glycoprotein glucosyltransferase. The EMBO Journal 14:4196–4203.

58. Labriola C, Cazzulo JJ, Parodi AJ (1999) Trypanosoma cruzi calreticulin is a lectin that binds monoglucosylated oligosaccharides but not protein moieties of glycoproteins. Mol Biol Cell 10:1381–1394.

59. Taylor SC, Thibault P, Tessier DC, et al (2003) Glycopeptide specificity of the secretory protein folding sensor UDP-glucose glycoprotein:glucosyltransferase. EMBO reports 4:405–411. doi: 10.1038/sj.embor.embor797

60. Wada I, Kai M, Imai S, et al (1997) Promotion of transferrin folding by cyclic interactions with calnexin and calreticulin. The EMBO Journal 16:5420–5432. doi: 10.1093/emboj/16.17.5420

61. Jin H, Yan Z, Nam KH, Li J (2007) Allele-specific suppression of a defective brassinosteroid receptor reveals a physiological role of UGGT in ER quality control. Molecular Cell 26:821–830. doi: 10.1016/j.molcel.2007.05.015

62. Pearse BR, Gabriel L, Wang N, Hebert DN (2008) A cell-based reglucosylation assay demonstrates the role of GT1 in the quality control of a maturing glycoprotein. J Cell Biol 181:309–320. doi: 10.1083/jcb.200712068

63. Li J, Zhao-Hui C, Batoux M, et al (2009) Specific ER quality control components required for biogenesis of the plant innate immune receptor EFR. Proc Natl Acad Sci USA 106:15973–15978. doi: 10.1073/pnas.0905532106

64. Pankow S, Bamberger C, Calzolari D, et al (2015) ΔF508 CFTR interactome remodelling promotes rescue of cystic fibrosis. Nature 528:510–516. doi: 10.1038/nature15729

65. Molinari M, Galli C, Vanoni O, et al (2005) Persistent glycoprotein misfolding activates the glucosidase II/UGT1-driven calnexin cycle to delay aggregation and loss of folding competence. Molecular Cell 20:503–512. doi: 10.1016/j.molcel.2005.09.027

