## Supplementary figures and images for "Clamping, bending, and twisting inter-domain motions in the misfold-recognising portion of UDP-glucose:glycoprotein glucosyl-transferase"

### SI Appendix movie I

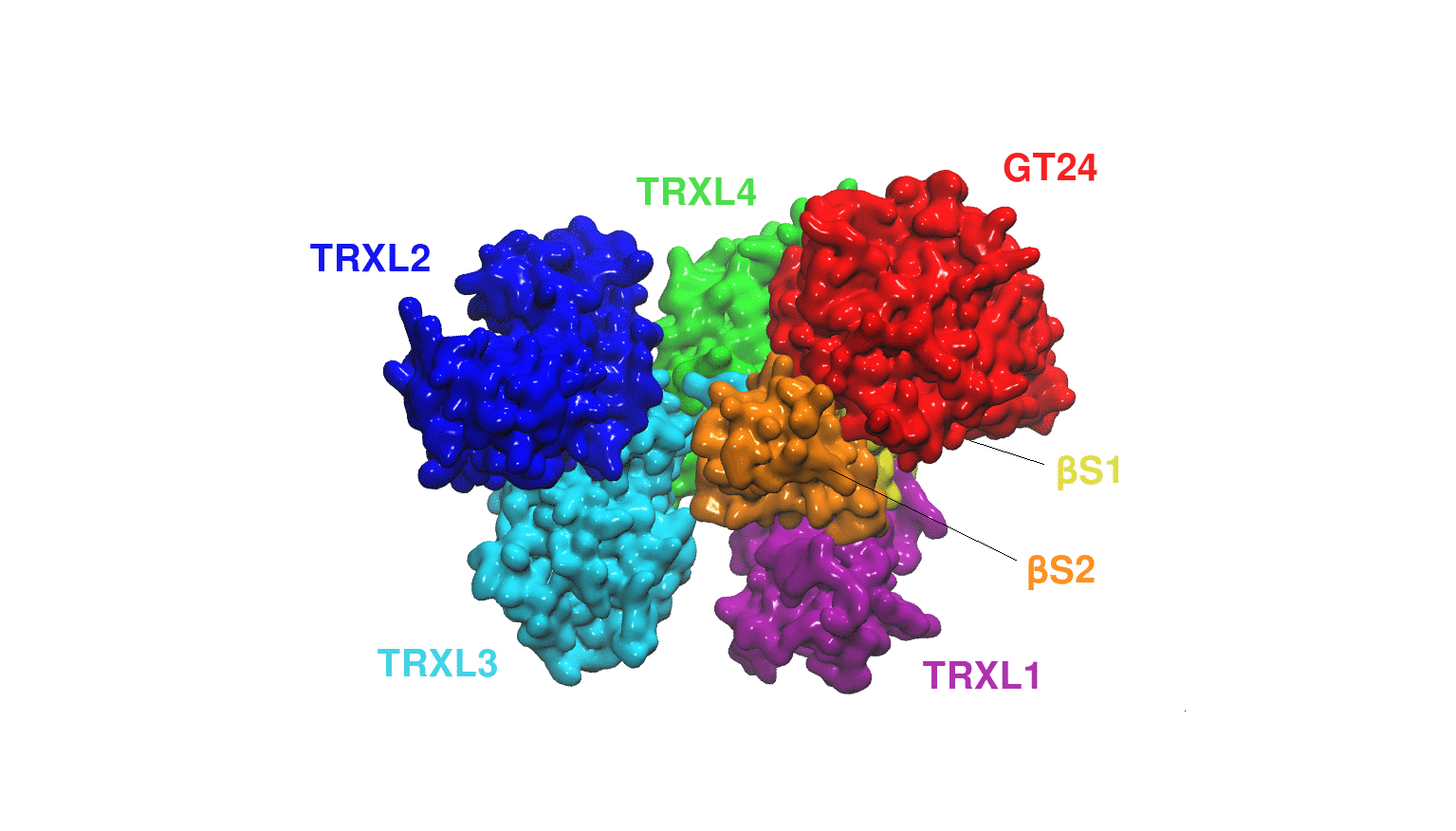

### SI Appendix movie II

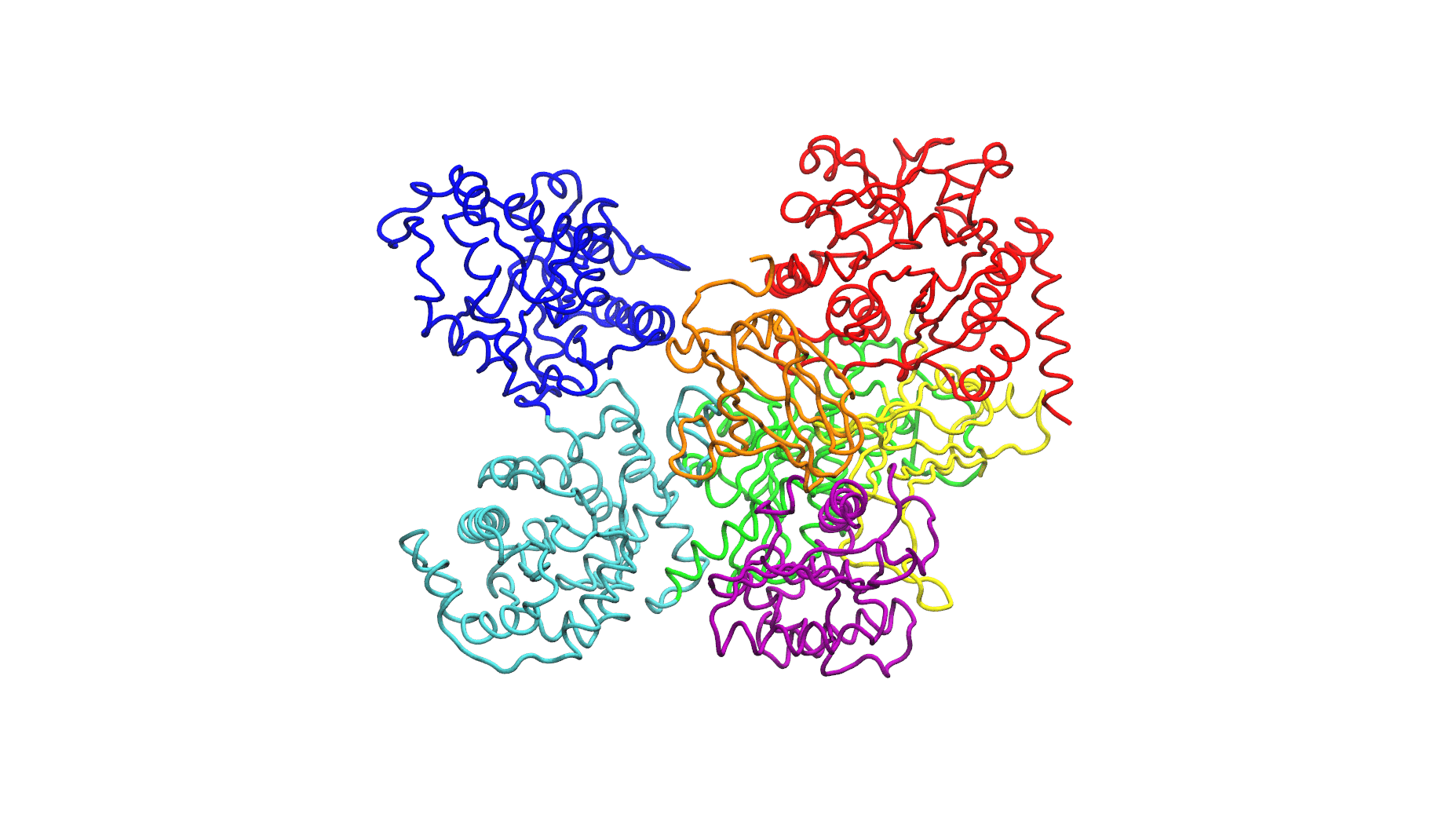

### SI Appendix movie III

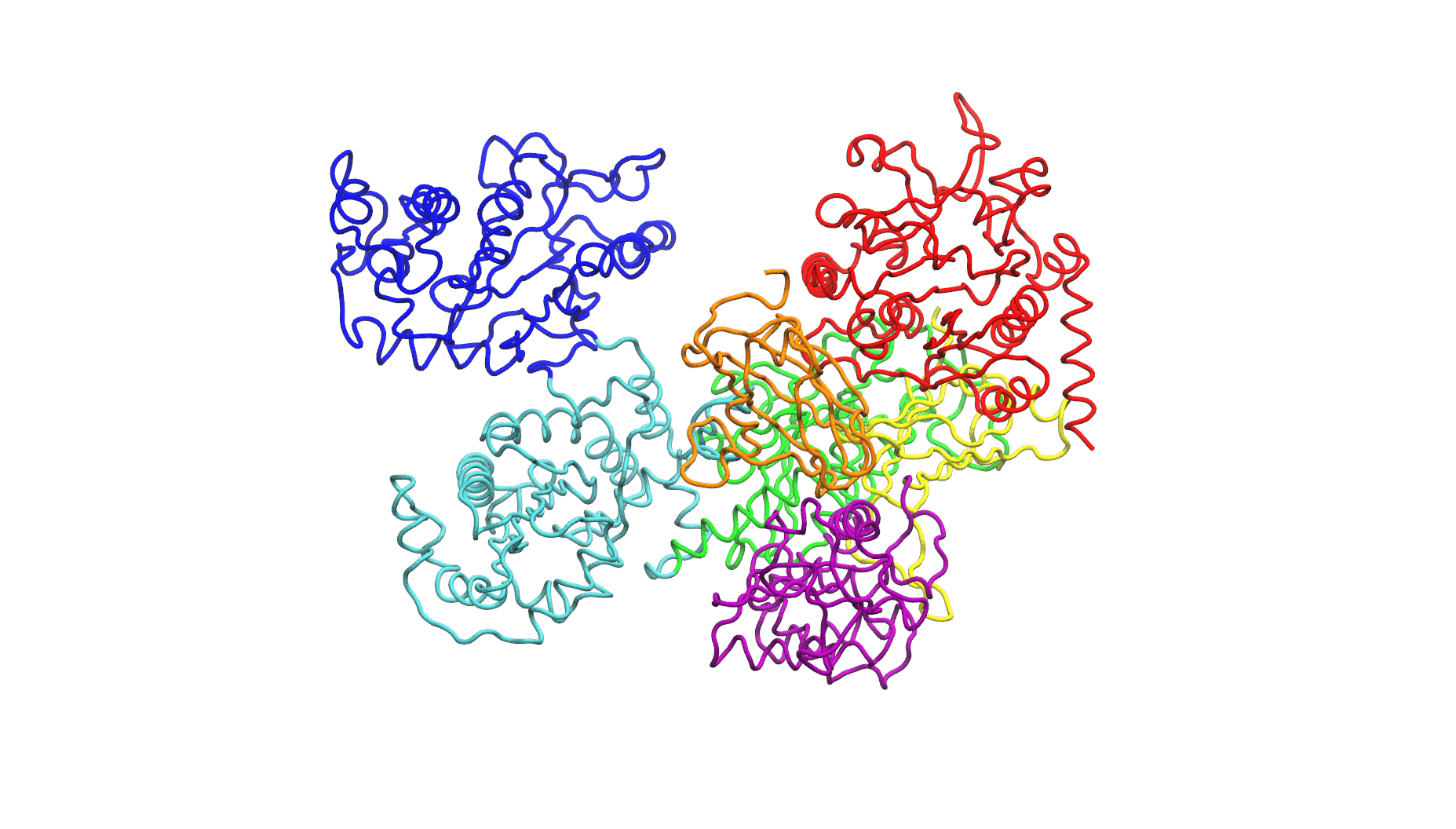

### SI Appendix movie IV

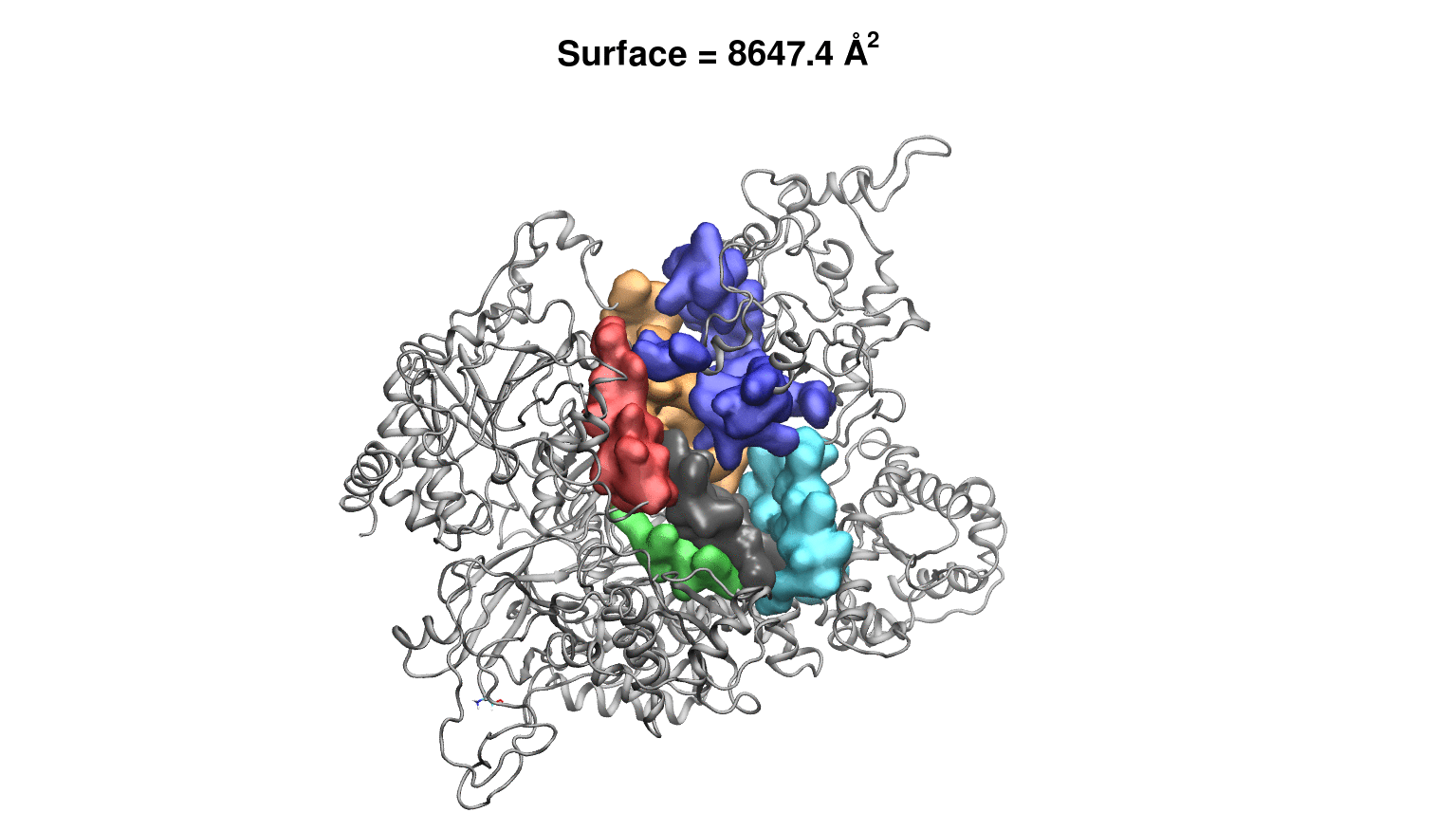
